# Dynamic profiling and functional interpretation of histone lysine crotonylation and lactylation during neural development

**DOI:** 10.1101/2021.05.21.444394

**Authors:** Shang-Kun Dai, Pei-Pei Liu, Zhao-Qian Teng, Chang-Mei Liu

## Abstract

**Background:** Metabolites such as crotonyl-CoA and lactyl-CoA influence gene expression through covalently modifying histones, known as histone lysine crotonylation (Kcr) and histone lysine lactylation (Kla). However, their existence patterns, dynamic changes, biological functions, as well as associations with histone lysine acetylation (Kac) and gene expression during mammalian development remain largely unknown, which imped us from understanding the epigenetic regulations of histone lysine acylations.

**Results:** Histone Kcr and Kla are widely distributed and undergo global changes during neural development. By profiling genome-wide dynamics of H3K9ac, H3K9cr and H3K18la in combination with ATAC sequencing and RNA sequencing, we reveal that these histone marks cooperatively regulate chromatin state and gene expression, as well as promote transcriptome remodelling to favour cell fate transition in the developing telencephalon. Importantly, we demonstrate that global histone Kcr and Kla are not consequence of transcription and furtherly identify histone deacetylase 1-3 (HDAC1-3) as novel “erasers” of H3K18la. Taking advantage of induced differentiation system of P19 embryonal carcinoma (EC) cells, we find that a selective inhibitor of HDAC1-3, MS-275 pre-activates neural fate-related transcriptional programs via stimulating multiple histone lysine acylations simultaneously.

**Conclusions:** Our study uncover the interplays between histone lysine acylations to regulate gene expression and the differentiation-promoting functions of histone Kcr and Kla during development, and provide evidence that multiple histone lysine actlations synchronously alternate to orchestrate transcriptome responding under HDACs inhibition.

## Background

The N-terminal tails of histones are subjected to various post-translational modifications (PTMs), which shape local chromatin landscape to regulate transcription, replication, DNA repair and higher-order chromatin structure [1, 2]. Some metabolites have recently been reported as donors for histone PTMs to influence gene expression, including propionyl-CoA, butyl-CoA, crotonyl-CoA, lactyl-CoA, 2-hydroxyisobutyl-CoA, β-hydroxybutyryl-CoA, malonyl-CoA, succinyl-CoA and glutaryl-CoA [1, 3]. Of these, crotonyl-CoA and lactyl-CoA, which act as substrates for histone lysine crotonylation (Kcr) and lysine lactylation (Kla) reactions catalysed by p300/CBP respectively [4, 5], have attracted increasing attention due to their emerging roles in linking cellular metabolism and epigenetic regulation to regulate gene expression, cell state transitions and diseases progression [5-8].

Kcr and Kla have been identified in both H2A, H2B, H3 and H4, and they share the most common modification sites with Kac [5, 9]. Over the last five years, regulatory mechanisms of histone Kcr have been thoroughly elucidated, including its “writers” (enzymes in charge of catalysing Kcr) [4, 10, 11], “erasers” (enzymes responsible for removing Kcr) [12-14], “readers” (enzymes capable of recognising Kcr) [15-18], and metabolic regulators (enzymes involving in crotonyl-CoA metabolism) [4, 19-23], which extend our knowledge about versatile functions of histone Kcr. Most recent studies have demonstrated that p300 and glycolysis are major pathways to regulate histone Kla levels, and suggested that H3K18la involves in M1 macrophage polarization, cell reprograming and tumorigenesis [5, 7, 8]. Both histone Kcr and Kla can directly stimulate transcription in a cell free system [4, 5]. Specifically, recognition of H3K9cr, H3K18cr and H3K27cr by YEATS domain of AF9 may regulate transcription process though recruiting the Super Elongation Complex (SEC) and Dot1L complex (DOT1LC) [15, 24, 25]. However, it is unclear whether the link between histone Kcr and Kla and transcription is causal or consequential in cells or in vivo under physiological conditions.

Genome-wide associations between histone Kcr and gene expression have been studied in various organisms [4, 9, 12, 14, 26-28], which indicate that it may act as a transcription activating histone PTMs. By contrast, histone Kla is only been reported to implicate in inflammation and oncogenesis-related transcription programs [5, 7]. However, it remains unknown whether and how histone Kcr and Kla involve in transcription regulation during development process. Given that histone Kac plays important roles in neurogenesis and neurological diseases [29-31], the potential functions of histone Kcr and Kla deserve to be explored during neural development. There is growing evidence that HDACs regulate multiple deacylation reactions except Kac [13, 32, 33], but we know little about how these histone lysine acylations are coordinated to regulate gene expression and influence biological process when HDACs are perturbed.

In the current study, we investigate the dynamic changes and associations with gene expression of histone Kcr and Kla, as well as interplays among histone lysine acylations during neural development. We observe the global existence and significant changes of histone Kcr and Kla in the developing telencephalon. Further multi-omics profiling uncovers that H3K9cr and H3K18la mark transcriptionally active chromatin state, but differ in signal pathways they involved both in vivo and in cells. Although there are distinct patterns of genome-wide alterations of H3K9ac, H3K9cr and H3K18la, in-depth comparisons of chromatin immunoprecipitation-sequencing (ChIP-seq) and RNA-seq data at different neural developmental stages reveal that these histone marks collaboratively regulate transcriptome remodelling to favour neural differentiation. In addition, inhibition of transcription dose not influence global histone Kcr and Kla levels. Finally, we discover that HDAC1-3 repression promotes neural differentiation-related transcriptional programs via multiple histone lysine acylations.

## Results

### Global changes of histone Kcr and Kla levels during neural development

We firstly determined the presence of several acylations modifications at sites of N-terminal tails of H3 in the embryonic telencephalon. The results demonstrated that these histone marks exhibited extensive existence in the germinal zones with strong staining signals of all antibodies (Fig. 1a, Additional file 1: Fig. S1a). Subsequently, we detected the levels of these histone marks across different developmental stages, and found that H3K9cr and H3K18cr levels increased but H3K18la and H3-Kla levels reduced at late-stage neurogenesis compared with early-stage neurogenesis, however, there were no changes of H3K14la and H3-Kcr levels (Fig. 1b, Additional file 1: Fig. S1b-d).

**Fig. 1.**
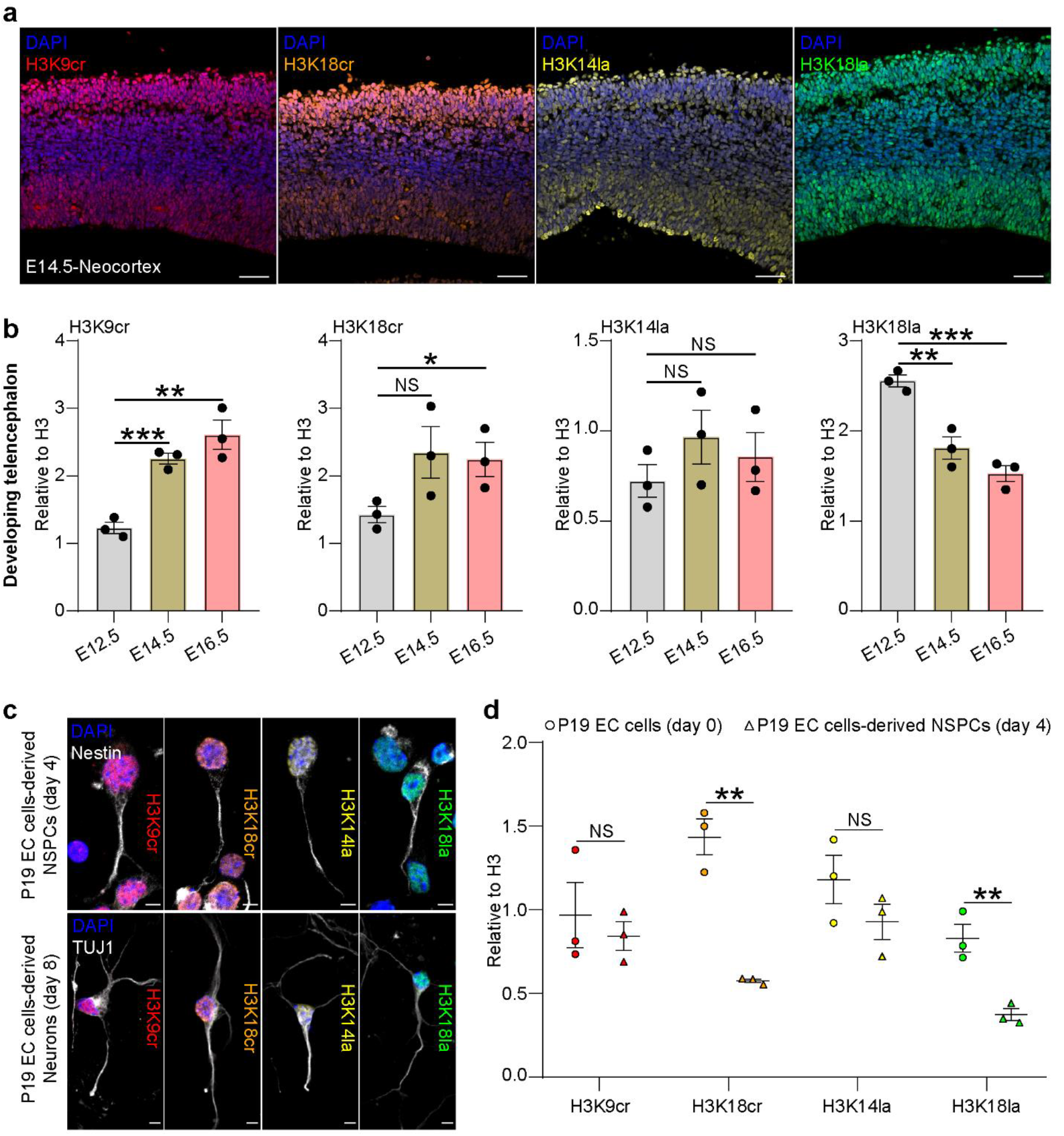
Existence and changes of histone Kcr and Kla during neural development. **a** Immunostaining of indicated histone marks on mouse forebrain sections at E14.5 (n = 1). Scale bar, 50 μm. **b** Statical analysis for changes of indicated histone marks in the developing telencephalon by western blot assay (n = 3). **c** Immunocytochemistry of indicated histone marks and markers for P19 EC cells-derived NSPCs (Nestin) or neurons (TUJ1) (n = 1). Scale bar, 5 μm. **d** Statical analysis for changes of indicated histone marks during induced neural differentiation of P19 EC cells (n = 3). In (**b, d**), data were represented as mean ± SEM; NS: no significance; *P< 0.05, **P< 0.01, ***P< 0.001 by two-tailed unpaired Student’s t-test.

We then checked the global changes of these histone marks at different stages of neural differentiation in cells (Additional file 1: Fig. S1e). The results showed that these histone marks could be observed in both Nestin^+^ neural stem/progenitor cells (NSPCs) and TUJ1^+^ neurons derived from P19 EC cells (Fig. 1c). Interestingly, the levels of H3K18cr and H3K18la, but not H3K9cr and H3K14la declined sharply during neural differentiation of P19 EC cells (Fig. 1d, Additional file 1: Fig. S1f), which indicated distinct dynamic changes of site-specific histone lysine acylations during cell fate transition. Altogether, these results suggest that histone Kcr and Kla are widely distributed and undergo global changes during neural development.

### Multi-omic profiling and functional interpretation of H3K9cr and H3K18la

In order to explore the chromatin function of histone Kcr and Kla during neural development, we performed specific anti-H3K9ac, H3K9cr and H3K18la (lactylation is not found at H3K9 site in mouse cells) ChIP-seq assays of the telencephalon at E13.5 (Additional file 1: Fig. S2) [5], and generated highly reproducible sequencing data (Additional file 1: Fig. S3a). The results suggested that the majority of H3K9ac, H3K9cr and H3K18la peaks were located at the promoter regions (Fig. 2a). Immediately, k-means clustering of the annotated genes revealed that gene loci with higher levels of H3K9ac were usually accompanied with higher levels of H3K9cr, H3K18la, chromatin accessibility as well as gene expression (Fig. 2b). H3K9ac-marked genes were also largely modified with H3K9cr and H3K18la (Additional file 1: Fig. S3b). Furthermore, genes with all three marks displayed the higher chromatin accessibility in the promoter regions and higher gene expression levels than that of the mono-marked genes (Fig. 2c). These results suggested that different types of histone lysine acylations might co-contribute to transcriptionally active chromatin state.

**Fig. 2.**
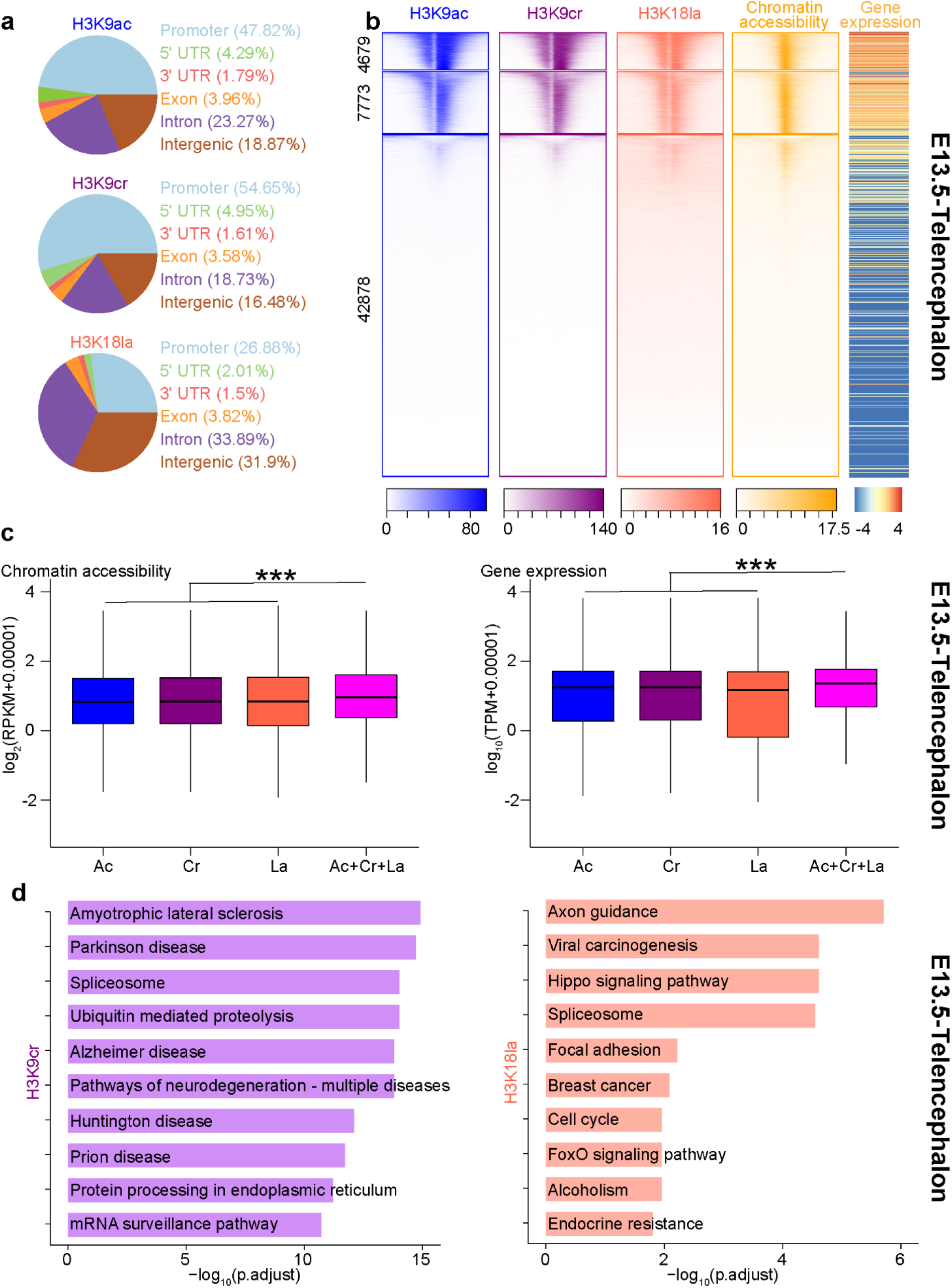
Multi-omics profiling and functional annotation of H3K9cr and H3K18la in vivo. **a** Pie chart showing distribution of peaks of indicated histone marks at annotated genomic regions in the E13.5 telencephalon. **b** Heatmaps of indicated histone marks intensity (scaled by RPKM), chromatin accessibility levels (scaled by RPKM) plotted within +/-2.5 kb of the TSS (5’ to 3’), and gene expression levels (scaled by log_10_(TPM+0.00001)) which were divided to into three clusters and ranked from the highest to the lowest by H3K9ac signal in the E13.5 telencephalon, the number of genes in each cluster was labelled on the left. **c** Boxplot showing comparison of changes of chromatin accessibility (left panel) and gene expression (right panel) across different groups of genes in the E13.5 telencephalon. Ac, Cr, La and Ac+Cr+La: genes under regulation of H3K9ac (13789 genes), H3K9cr (13726 genes), H3K18la (13113 genes) or all three marks (10162 genes). **d** KEGG pathway enrichment analysis of genes with the strongest H3K9cr signal (left panel, 3561 genes) and H3K18la signal (right panel, 3568 genes) in the E13.5 telencephalon. In (**c**), ***P< 0.001 by two-tailed unpaired Student’s t-test.

In our analysis, we found that there were many genes marked by these histone marks with different levels of modification (Additional file 1: Fig. S3b, c). Therefore, we divided peaks into three clusters based on their modification levels to interpret biological functions of these histone lysine acylations (Additional file 1: Fig. S3c). Interestingly, genes with the strongest H3K9cr intensity (cluster 1) were annotated in RNA and protein metabolism and neurodegenerative disorders-related KEGG pathways, however, genes with the highest H3K18la levels were mainly involved in cell proliferation-related pathways (Fig. 2d). Consistently, our ChIP-seq assays of these histone marks using P19 EC cells-derived NSPCs confirmed their interplays and important functions at the cell level (Additional file 1: Fig. S4). Thus, these results indicate that H3K9ac, H3K9cr and H3K18la exhibit cooperative regulation of chromatin state and gene expression, but they are involved in different signalling pathways in vivo and in cells.

### Genome-wide alterations in H3K9cr and H3K18la regulate transcriptome remodelling to favour neural differentiation in vivo

Histone marks undergo dynamic changes and play important roles during embryonic neurogenesis in which NSPCs exit cell cycle and differentiate into neurons [31]. However, it is still not clear how histone Kcr and Kla participate in this process. Therefore, we compared genome-wide changes of H3K9ac, H3K9cr and H3K18la at early (E13.5) and late (E16.5) stages of embryonic neurogenesis (Additional file 1: Fig. S5). We first explored the association between alterations of these histone marks in the promoter regions and gene expression changes (Additional file 1: Fig. S6a, b; Additional file 2: Table S1), and found that genes with different binding of all three marks displayed the higher gene expression changes than genes with alterations of mono-mark (Fig. 3a, b). Importantly, there were significant correlations between H3K9ac, H3K9cr and H3K18la enrichment and altered gene expression (Additional file 1: Fig. S6c-e). After checking genes with different patterns of changes of these histone marks, we found that H3K18la enrichment declined more significantly at genes encoding histones which play important roles during cell proliferation than that of H3K9ac and H3K9cr (Additional file 1: Fig. S6f, g) [34].

**Fig. 3.**
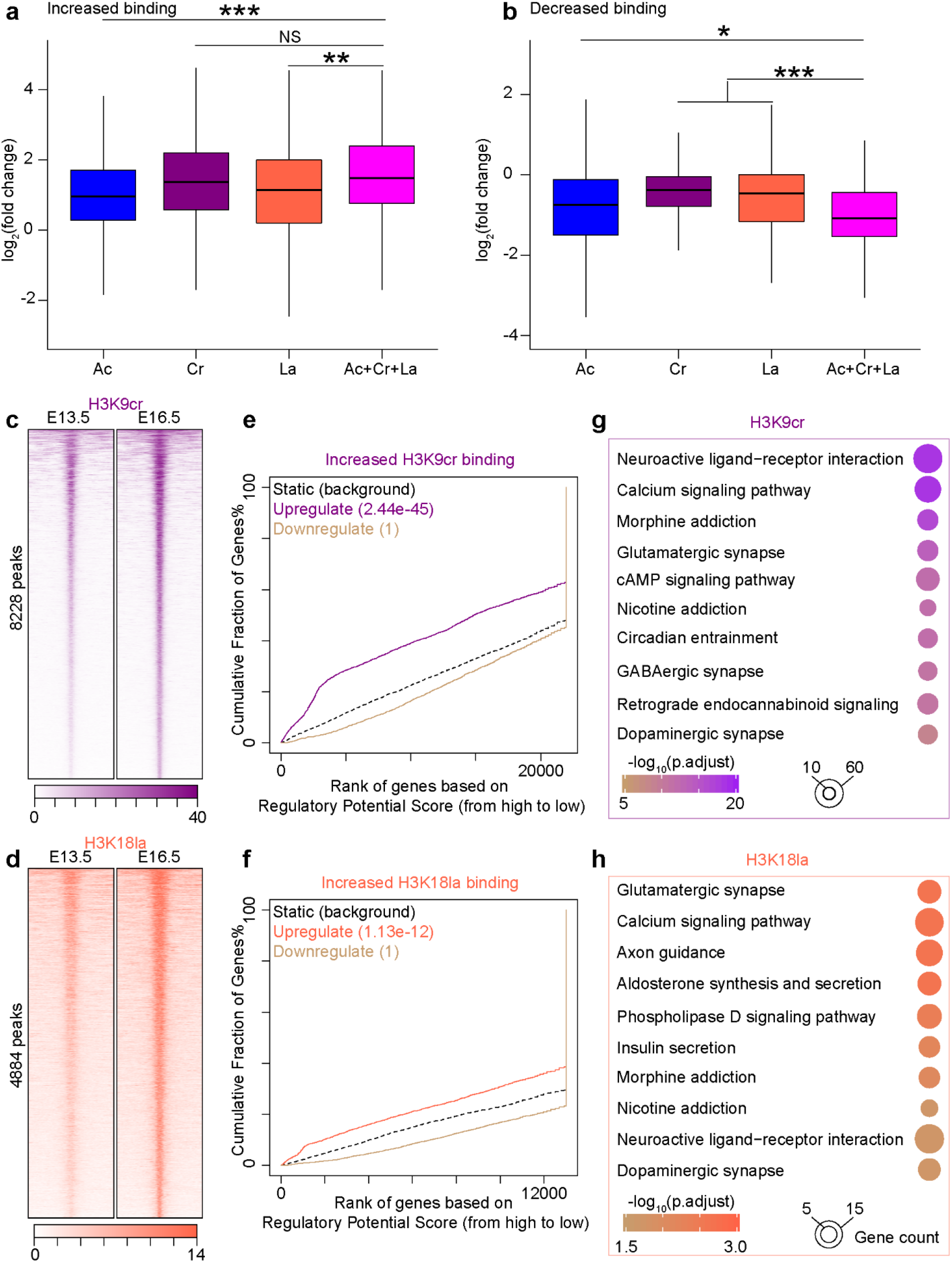
Genome-wide changes of H3K9cr and H3K18la during neural development. **a** Boxplot showing comparison of gene expression changes with increased binding of indicated histone marks in the promoter regions across different groups of genes. Ac, Cr, La and Ac+Cr+La: genes with increased binding of H3K9ac (3717 genes), H3K9cr (1785 genes), H3K18la (542 genes) or all three marks (196 genes). **b** Boxplot showing comparison of gene expression changes with decreased binding of indicated histone marks in the promoter regions across different groups of genes. Ac, Cr, La and Ac+Cr+La: genes with decreased binding of H3K9ac (784 genes), H3K9cr (2693 genes), H3K18la (730 genes) or all three marks (178 genes). **c, d** Heatmaps at peaks with increased H3K9cr (**c**) or H3K18la (**d**) binding (scaled by RPKM) plotted within +/-5 kb of the peak center (5’ to 3’), the number of peaks was labelled on the left. **e, f** BETA plot of combined computational analysis of ChIP-seq and RNA-seq data using peaks with increased H3K9cr (**e**) or H3K18cr (**f**) binding. **g, h** KEGG pathway enrichment analysis of up-regulated genes with increased H3K9cr (**g**) or H3K18la binding (**h**), there were 923 and 1342 genes in (**e**) and (**f**) respectively. In (**a, b**), NS: no significance; *P< 0.05, **P< 0.01, ***P< 0.001 by two-tailed unpaired Student’s t-test.

Next, we compared the peaks with different binding of H3K9ac, H3K9cr and H3K18la during neural development. Unexpectedly, changes in number of peaks and annotated genes with different binding of modifications showed distinct patterns across these histone marks (Additional file 1: Fig. S7, Additional file 3: Table S2). For instance, H3K9ac was the mark with the most peaks and annotated genes with increased enrichment, and tailed the fewest with reduced binding comparing with H3K9cr and H3K18la. In contrast, there were more decreased H3K18la binding than increased as either the number of peaks or annotated genes versus H3K9ac and H3K9cr (Additional file 1: Fig. S7a-e). However, involved biological processes of genes with increased or decreased binding of all three marks were highly consistent with development program (Additional file 1: Fig. S5, S7f and g). To correlate the alterations of these histone marks with gene expression changes, we performed binding and expression target analysis (BETA) [35], and revealed that H3K9cr and H3K18la peaks with increased enrichment had significant effects as a gene activator, while peaks of reduced H3K9cr and H3K18la binding could act as the predictor of down-regulated gene expression (Fig. 3c-f, Additional file 1: Fig. S8a-d). Further enrichment analysis showed that up-regulated genes with increased H3K9cr or H3K18la enrichment were mainly involved in neural differentiation (Fig. 3g, h), comparatively, while down-regulated genes with reduced binding of H3K9cr or H3K18la were tightly associated with cell proliferation (Additional file 1: Fig. S8e, f). Collectively, although there is a huge difference in the number of peaks and annotated genes with different binding across H3K9ac, H3K9cr and H3K18la modifications, these histone marks undergo genome-wide alterations and collaborate to regulate gene expression associated with neural differentiation in vivo.

### Transcription repression has no effect on global histone Kcr and Kla levels

A recent study shows that inhibition of transcription results in the loss of histone Kac in bulk histones, and majority of histone acetylation is dependent on transcription [36]. However, the associations between non-acetylation histone lysine acylations and transcription under physiological conditions are unclear. To investigate if the histone Kcr and Kla were influenced by transcription, we treated P19 EC cells-derived NSPCs (day 2) with the well-studied transcription inhibitors Triptolide (inhibiting transcription initiation) and Flavopiridol (inhibiting transcription elongation) at high concentrations [37, 38]. Transcription was significantly inhibited as indicated by the loss of RNA polymerase II (RNAP2) serine 5 CTD phosphorylation (S5p) and RNAP2 serine 2 CTD phosphorylation (S2p) (Fig. 4a, b). Surprisingly, we didn’t detect any significant changes of H3K9cr, H3K18cr, H3K14la and H3K18la levels (Fig. 4c-e), which indicate that majority of histone Kcr and Kla are not consequence of transcription.

**Fig. 4.**
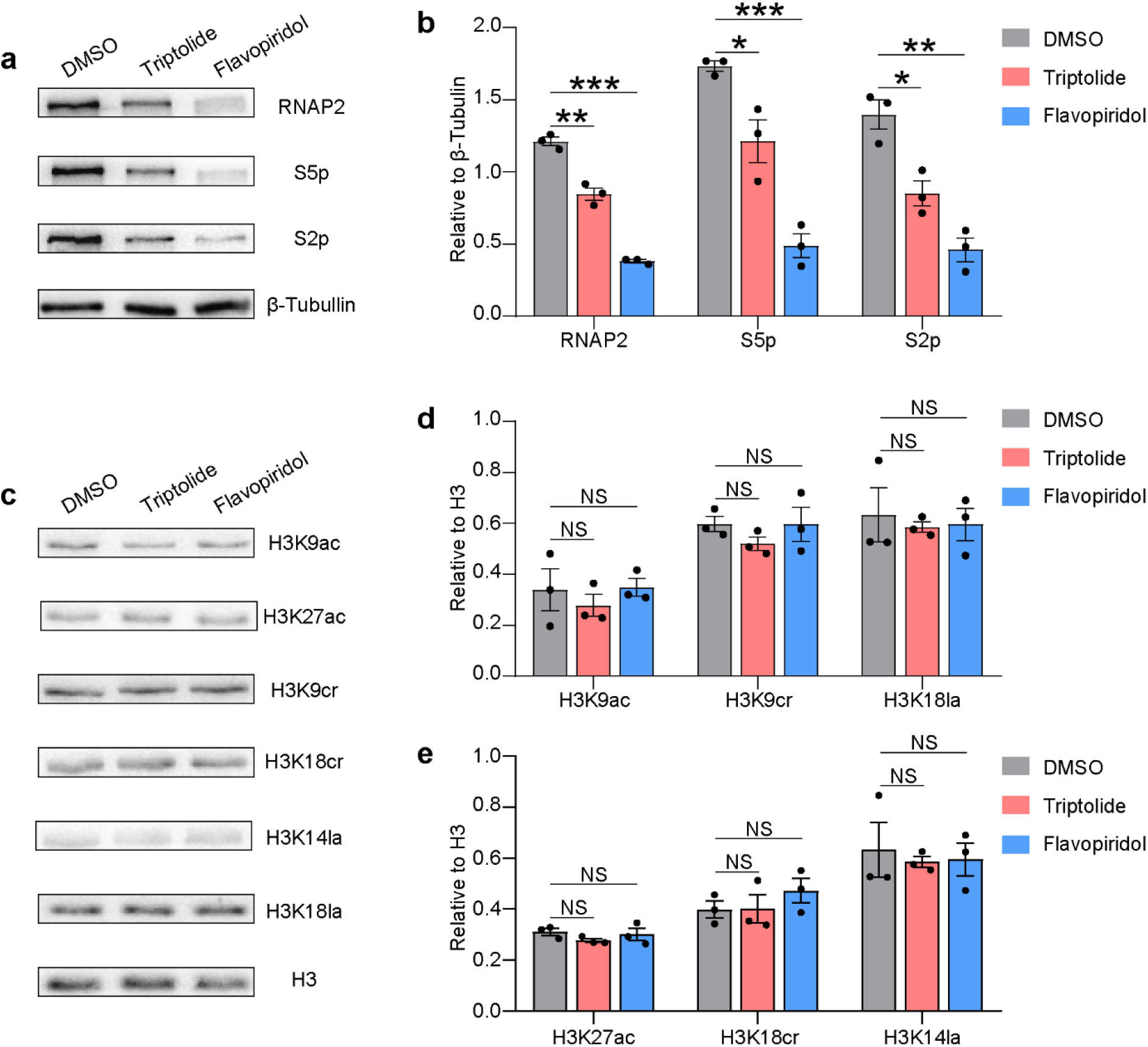
The association between transcription inhibition and histone Kcr and Kla levels. **a, b** Representative plots (**a**) and quantification analysis (**b**) of changes of RNA polymerase II (RNAP2), RNAP2 serine 5 CTD phosphorylation (S5p) and RNAP2 serine 2 CTD phosphorylation (S2p) levels in P19 EC cells-derived NSPCs treated with DMSO, Triptolide or Flavopiridol (n = 3). **c-e** Representative plots (**c**) and quantification analysis (**d, e**) of changes of levels of indicated histone marks in P19 EC cells-derived NSPCs treated with DMSO, Triptolide or Flavopiridol (n = 3). In (**b, d, e**), data are represented as mean ± SEM ; NS: no significance; ^*^P< 0.05, ^**^P< 0.01, ^***^P< 0.001 by two-tailed unpaired Student’s t-test.

### HDAC1-3 inhibition stimulates the elevation of H3K18la both in cells and in vivo

Given the fact that some HDACs have been demonstrated to “erase” various histone lysine acylations except Kac [13, 32], we next asked whether HDACs could regulate histone Kla. To this end, we applied two pan-HDACs inhibitors (SAHA and VPA), and found that SAHA treatment increased H3K18la levels while VPA stimulated both H3K14la and H3K18la in P19 EC cells (Fig. 5a, b). Moreover, we found that a selective inhibitor of HDAC1-3, MS-275, robustly promoted the elevation of H3K14la and H3K18la comparing with EX 527 (a selective SIRT1 inhibitor) (Fig. 5c). Meanwhile, we observed enhanced H3K9cr and H3K18cr levels after SAHA, VPA or MS-275 treatment, which were consistent with previous studies (Fig. 5a-c) [12].

**Fig. 5.**
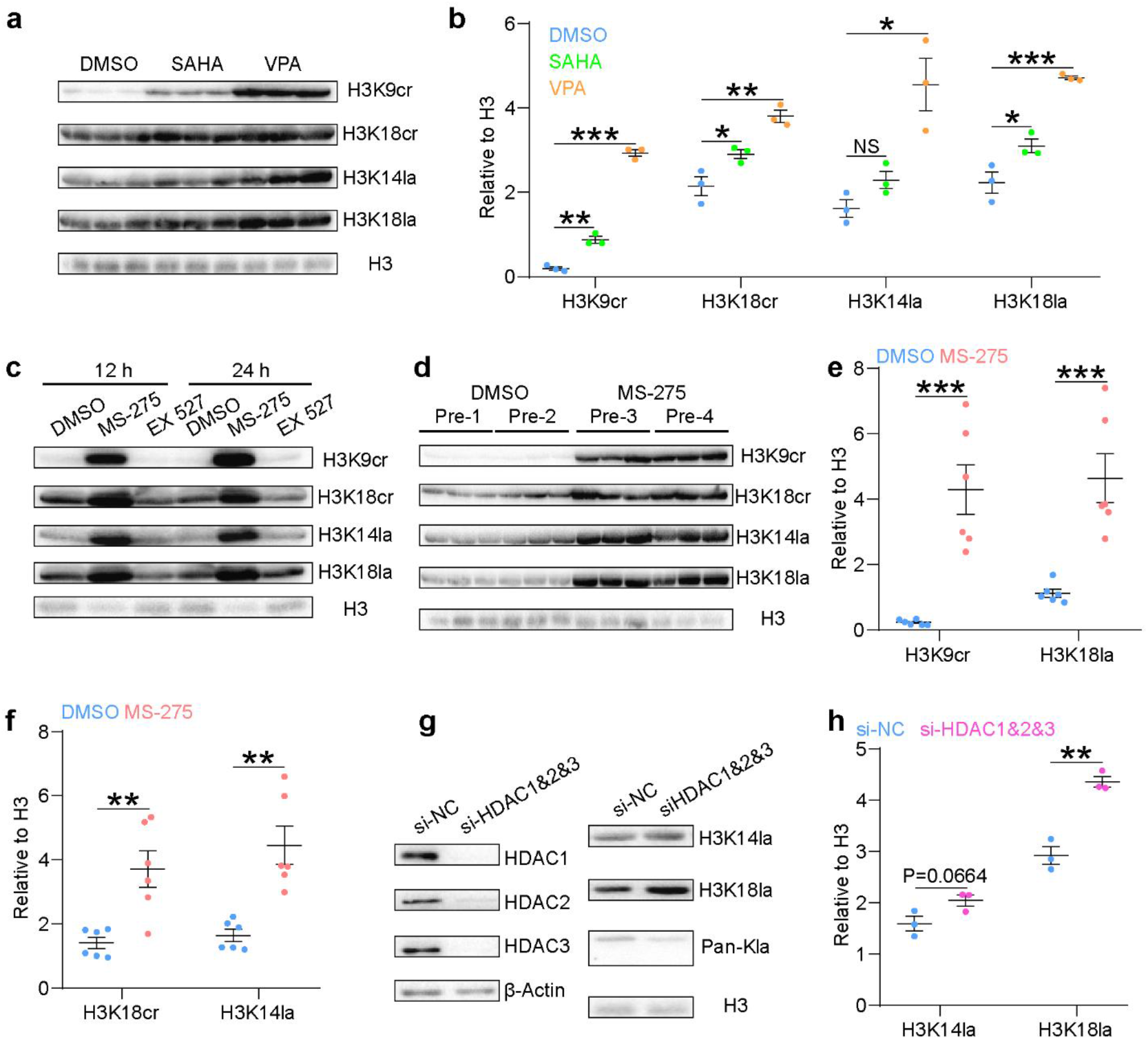
HDAC1-3 inhibition stimulates histone Kcr and Kla levels. **a, b** Representative plots (**a**) and quantification analysis (**b**) of changes of indicated histone marks in DMSO, SAHA or VPA treated P19 EC cells (n = 3). **c** Western blot assay of changes of indicated histone marks in DMSO, MS-275 or EX 527 treated P19 EC cells for different time intervals (n = 1). **d-f** Western blot assay (**d**) and quantification analysis (**e, f**) of changes of indicated histone marks in the telencephalon at E15.5 injected with DMSO or MS-275 for two consecutive days (E13.5-E15.5), Pre: pregnant mice. **g** Representative plots showing efficient knockdown of *Hdac1-3* (left panel) and changes of histone Kla levels (right panel) in P19 EC cells. **h** Quantification analysis of changes of H3K14la and H3K18la levels after *Hdac*1-3 knockdown in P19 EC cells (n = 3). In (**b, e, f, h**), data are represented as mean ± SEM ; NS: no significance; *P< 0.05, **P< 0.01, ***P< 0.001 by two-tailed unpaired Student’s t-test.

To confirm our above findings in vivo, we injected pregnant mice with MS-275 and detected significant increases in H3K14la and H3K18la levels, as well as an enhancement of H3K9cr and H3K18cr levels which have not been reported previously in vivo (Fig. 5d-f). We next investigated whether the dynamic changes of histone Kla existed after interference of expression of HDACs, the results showed that simultaneous knockdown of *Hdac1-3* increased levels of H3K18la but not global histone Kla in P19 EC cells (Fig. 5g, h), which might be caused by different regulatory mechanisms at other histone Kla sites. Overall, these results suggest that HDAC1-3 are novel site-specific histone lysine delactylases both in cells and in vivo.

### Multiple histone lysine acylations are implicated in the upregulation of neural differentiation-related genes by MS-275

H3K18la has been demonstrated to regulate M1 macrophage polarization, cell reprograming and tumorigenesis recently [5, 7, 8], however, we still don’t know how H3K18la is regulated and co-operates with other histone lysine acylations during neural cell fate determination. To this end, we treated differentiating P19 EC cells with MS-275, and performed ChIP-seq assays of H3K9ac, H3K9cr and H3K18la subsequently. HDAC1-3 inhibition by MS-275 induced genome-wide changes of H3K9ac, H3K9cr and H3K18la binding (Additional file 4: Table S3, Additional file 1: Fig. S9a-c). Interestingly, the enhanced enrichment of these histone marks both promoted the expression of neural differentiation-related genes (Additional file 1: Fig. S9d-i). Furtherly, genes with increased binding of all three marks were tightly associated with neural cell fate decisions (Fig. 6a, b; Additional file 1: Fig. S10a), and displayed the higher gene expression changes than those with alterations of mono-mark after MS-275 treatment in differentiating P19 EC cells (Fig. 6c).

**Fig. 6.**
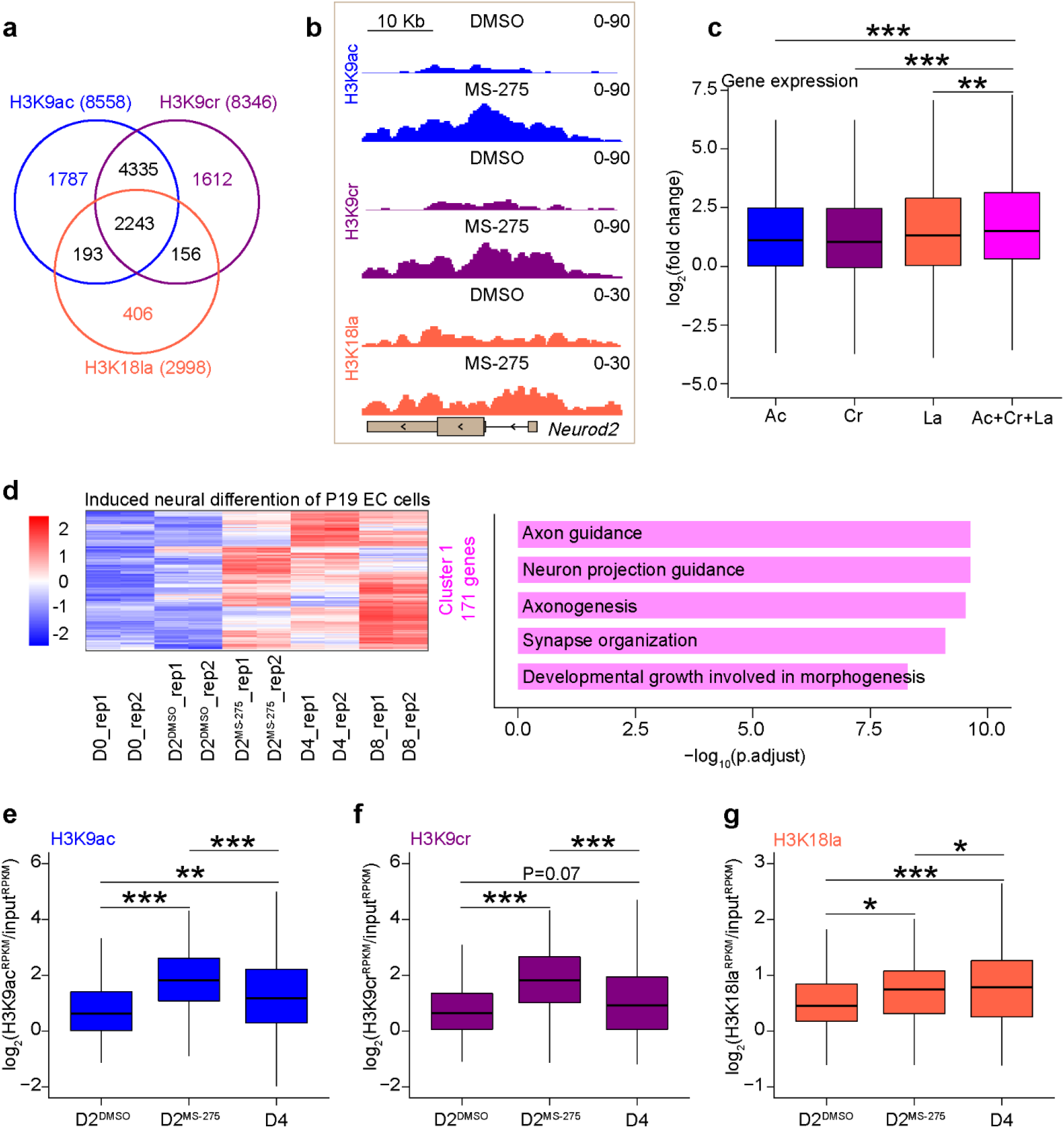
HDAC1-3 regulates neural cell fate determination via multiple histone lysine acylations. **a** Venn diagram of a combined comparison of groups of genes under regulation of H3K9ac, H3K9cr or H3K18la, with increased binding of each mark. **b** Representative genome-browser view showing increased multiple histone lysine acylations at *Neuord2* gene of different sequencing data sets after MS-275 treatment in P19 EC cells-derived NSPCs. **c** Boxplot showing comparison of gene expression changes with increased indicated histone marks binding in the promoter regions across different groups of genes. Ac, Cr, La and Ac+Cr+La: genes with increased binding of H3K9ac (4883 genes), H3K9cr (4757 genes), H3K18la (1197 genes) or all three marks (744 genes). **d** Heatmap showing dynamic changes of up-regulated genes (DESeq2 vsd-normalized RNA-seq counts and scaled by row) with increased binding of all three marks (H3K9ac, H3K9cr and H3K18la) after MS-275 treatment during neural differentiation process of P19 EC cells, and these genes were divided into three clusters (see also Additional file 1: Fig. S10c) according their expression patterns (left panel), with the results of GO enrichment analysis (biological process) of genes in each cluster on right panel. **e-g** Boxplot showing comparison of changes of H3K9ac (**e**), H3K9cr (**f**) or H3K18 (**g**) levels across different groups of NSPCs at genes in cluster 1 (171 genes) defined in (**d**), which indicated that genes in cluster 1 got increased binding of these histone marks in MS-275 treated NSPCs at Day2 (D2^MS-275^) comparing with that in DMSO treated control group (D2^DMSO^), and the increased trends were also observed during normal neural differentiation (D4 (NSPCs at Day4) vs D2^DMSO^). In (**c, e-g**), *P< 0.05, **P< 0.01, ***P< 0.001 by two-tailed unpaired Student’s t-test.

Having taken gene expression changes into account, there were 635 up-regulated genes with increased binding of all three marks (Additional file 1: Fig. S10b), which could be divided into three clusters based on their patterns of expression changes across different timepoints of differentiation (Fig. 6d, Additional file 1: Fig. S10c). Cluster 1 containing genes with gradual upregulation of expression during differentiation, but pre-activated via advanced enhancement of multiple histone lysine acylations under HDAC1-3 inhibition by MS-275 (Fig. 6d-g). Conversely, genes in cluster 2 might be associated with the development of other germ layers, because both expression and histone lysine acylations levels of genes this cluster were unchanged during differentiation (Additional file 1: Fig. S10c-f). In addition, Genes in cluster 3 were highly expressed in P19 EC cells but down-regulated during differentiation, and they were involved in maintaining identity of P19 EC cells. Overall, MS-275 promotes transcriptional programs associated with neural differentiation in P19 EC cells via activating multiple histone lysine acylations simultaneously.

## Discussion

In this study, we have profiled dynamic changes and performed functional interpretation of H3K9cr and H3K18la in the developing telencephalon. We also demonstrate that HDAC1-3 act as novel “erasers” of H3K18la and furtherly link multiple histone lysine acylations with neural differentiation. First, these histone marks display extensive existence and significant changes during neural development. Next, both H3K9ac, H3K9cr and H3K18la are histone PTMs marking active chromatin regions and cooperatively regulate gene expression. Comparison of alterations of these marks across different developmental stages reveal that genome-wide increased and decreased H3K9cr and H3K18la binding mainly involve in neural differentiation and cell proliferation processes respectively, highlighting the remodelling of these marks to orchestrate changes of gene expression and cell fate (Fig. 7a). Moreover, we show that global levels of histone Kcr an Kla are not influenced by transcription inhibition. Importantly, we uncover that HDAC1-3 can “erase” H3K18la in cells and in vivo, and regulate neural differentiation of P19 EC cells via multiple histone lysine acylations (Fig. 7b).

**Fig. 7.**
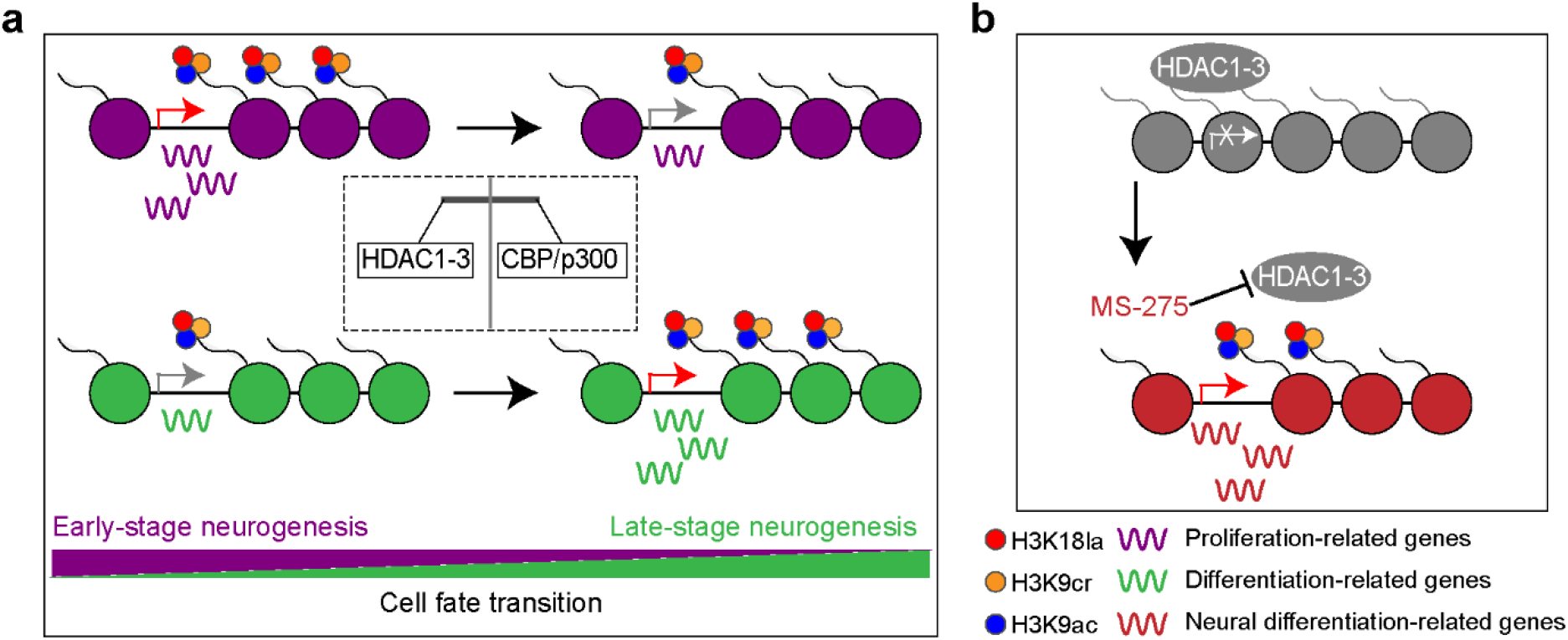
Working models of multiple histone lysine acylations to regulate gene expression during normal development and perturbed neural differentiation processes.

The extensive existence and so many marked-genes of histone Kcr and Kla in the developing telencephalon indicated that histone lysine acylations should act as universal epigenetic mechanisms to regulate gene expression during development. The difference in patterns of changes of H3K9ac, H3K9cr and H3K18la binding is understandable, because the levels of donors for various histone lysine acylations may alter during neural development. For instance, there is a transition from glycolytic to mitochondrial oxidative phosphorylation during neurogenesis, therefore it is possible that reduction in production of lactate and lactyl-CoA may influence dynamic changes of H3K18la and H3-Kla levels during development [39]. Knockdown of *CBP* inhibits neurogenesis in embryonic neocortex, and intermediate progenitors are prone to neural differentiation with *Hdac1* and *Hdac2* deletion in forebrain [29, 30], which indicate an opposite effect of these enzymes during neural differentiation. Both CBP and H3K9/14ac are already bind the promoter of *Tuj1* (a marker of neuron) at E13, binding peaks at E16, and then decrease postnatally in the developing cortex, correlating with neurogenesis [29]. In adult brain, CBP/p300 maintain the excitatory neuron identity though the regulation of histone Kac at cell type-specific promoter and enhancer regions [40]. These results suggest important roles of CBP/p300 during establishment and maintenance of neuron identity. At early stage of neurogenesis, neural differentiation-related genes are kept in primed state, and both histone acetyltransferases (HATs) and HDACs are targeted to these development-related primed genes simultaneously [41, 42]. Based on our findings that increased H3K9ac, H3K9cr and H3K18la binding at neural differentiation-related genes accompanied by undergoing neurogenesis and HDAC1-3 inhibition in differentiating P19 EC cells, as well as the capacity of CBP/p300 to catalyse histone Kac, Kcr and Kla [4, 10], we propose the existence of a switch of “actyl-spray” targeting regulatory elements of genes related to neural differentiation, which is triggered by the balance between CBP/p300 and HDACs (Fig. 7a).

Almost all studies of considering the causal relationship between histone Kcr and Kla and transcription or gene expression, make their conclusions based mainly on the consistency between dynamic changes of histone marks and gene expression. However, there is no evidence to support direct regulation of transcription by histone Kcr and Kla at the cell level. In addition, whether the link between histone Kcr and Kla and transcription under physiological condition is a causal or consequential event has not been addressed previously. Although the present study can’t rule out the possibility of changes of these histone marks in the local sites of the genome, our data clearly show that transcription inhibition has no effect on global histone Kcr and Kla levels, indicating that the majority of histone Kcr and Kla are not dependent on transcription. In support of our conclusion, recent studies report that there is no global effect on histone Kac within a short time frame of transcription inhibition [43].

Recently, some HDACs have been proved to “erase” histone Kcr (HDAC1, 2, 3, 8 and SIRT1) [13], histone lysine benzoylation (Kbz, SIRT2) [32] and histone lysine β-hydroxybutyrylation (Kbhb, HDAC1 and 2) [33] in cells. Surprisingly, we find that inhibition of HDAC1-3 significantly stimulates H3K14la and H3K18la levels both in cells and in vivo, which suggest that HDAC1-3 are novel “erasers” of histone Kla. A recent study by Olsen and collaborators also report histone delactylation activity of HDAC1-3 in vitro [44]. But under physiological conditions, they find that there are no changes in H3K18la levels upon knockdown of *HADC1-3* in HeLa cells. In contrast, we observe remarkably increased H3K18la levels after simultaneous knockdown of *Hdac1-3* in P19 EC cells. We speculate that this discrepancy may be due to the knockdown efficiency of HDAC1-3 and the cell types. Indeed, our in vivo and ChIP-seq data do support the conclusion that HDAC1-3 can “erase” H3K18la under physiological conditions. Functionally, genes with gain of H3K18la after MS-275 treatment are directly involved in neural differentiation (Additional file 1: Fig. S9c, f, i). Although there are two studies showing genome-wide changes of H3K18ac and H3K18cr under perturbation of HDACs, but the associations among histone lysine acylations and underlying biological functions are less studied [12, 14]. The highlight of our research is that we link HDACs and multiple histone lysine acylations with cell fate transition associated transcriptome program (Fig. 7b).

## Conclusions

Overall, our study provides the first and novel insights into the dynamic changes, epigenetic interactions, transcription regulation and biological functions of histone lysine acylations during neural development. We also bridge HDACs, multiple histone lysine acylations with neural differentiation, which highlight the cooperation and functional importance and of these histone marks. Our results provide new perspective for epigenetic regulation in development and diseases processes.

## Methods

### Mice

All mice used in the current study had a ICR (CD-1) background, pregnant mice were purchased from SiPeiFu Biotechnology Co., Ltd (Beijing, China). All mouse experiments were approved by the Animal Committee of the Institute of Zoology, Chinese Academy of Sciences, Beijing, China.

### Cell culture

P19 embryonal carcinoma (EC) cells were maintained in the proliferation medium (minimum essential medium-alpha (MEMα; Thermo Fisher Scientific) containing 10% fetal bovine serum (FBS; Thermo Fisher Scientific)) in an incubator at 37°C and 5% CO_2_. For induced neural differentiation, 1×10^6^ cells were cultured on a 9.6 cm bacterial-grade Petri dish (Greiner) in the induction medium (proliferation medium containing 1 μM all-trans retinoic acid (Sigma-Aldrich)) for 4 d, with media exchanged every 2 d. To induce neuronal differentiation, the aggregated neurospheres were digested in to single cells by Accutase (STEMCELL Technologies) and plated at a density of 1.2×10^7^ cells per 10 cm culture dish (Corning) pre-coated with 0.1 mg/ml poly-D-lysine (PDL; Sigma-Aldrich) in the differentiation medium (DMEM/F12 containing with 2% B27 (Thermo Fisher Scientific)) for 4 d, with media exchanged every 2 d.

### Inhibitor treatment

Inhibitors were all purchased from Selleck Chemicals. P19 EC cells treatments were performed in the proliferation medium at the following concentration for 12 h or 24 h: 0.5 μM SAHA (S1047, CAS 149647-78-9), 1 mM VPA (S3944, CAS:99-66-1), 1 μM MS-275 (S1053, CAS 209783-80-2) and 1 μM EX 527 (S1541, CAS 49843-98-3). For in vivo experiment, 20 mg/kg MS-275 was injected every 24 h for two consecutive days on pregnant mice at day 13.5 of pregnancy (E13.5-E15.5). P19 EC cells-derived NSPCs (day 1) were treated with 1 μM MS-275 for 1 d and then collected for ChIP-seq and RNA-seq assays. P19 EC cells-derived NSPCs (day 2) treatments were performed in the induced medium at the follow concentration for 1 h: 5 μM Triptolide (S3604, CAS 38748-32-2) and 5 μM Flavopiridol HCl (S2679, CAS 131740-09-5).

### Cell transfection

P19 EC cells were transfected with double-strand siRNA using Lipofectamine™ RNAiMAX (Thermo Fisher Scientific) with the final concentration at 50 nM for 3 d and then collected for western blot assay. The Negative control siRNA (siR NC #1, siN0000001-1-5) and siRNA targeting *Hdac1* (si-m-Hdac1_004, siB140729172936), *Hdac2* (si-m-Hdac2_004, siB140729171225) and *Hdac3* (si-m-Hdac3_001, siG141217112523) were purchased from Ribobio Co., Ltd (Guangzhou, China).

### Immunocytochemistry and immunostaining

For immunocytochemistry analyses, cells on coverslips were fixed in 4% paraformaldehyde for 15 min at room temperature, washed three times in PBS (10-min wash each time). Next, cells were blocked in a blocking solution (2% bovine serum albumin and 0.3% Triton X-100) at room temperature for 1 h. The coverslips were then incubated with primary antibodies diluted in blocking solution at 4°C overnight, and labelled using appropriate secondary antibodies at room temperature for 2 h. For immunostaining, embryonic mouse brains were cut into 12-μm-thick cryosections using Leica CM 1950. The staining procedures for brain slices were similar with those used for immunocytochemistry described above. All images were acquired by confocal microscopy (ZEISS, LSM880) and analysed with ZEN 2012. The detailed information of the primary and secondary antibodies used were summarized in the Additional file 1: Table S4.

### Western blot and dot blot assay

Total proteins were extracted using the RIPA lysis buffer (Beyotime Biotechnology, Shanghai, China) containing 1×complete proteinase inhibitor (Bimake), and protein concentration was defined using BCA protein assay kit (Beyotime). Histones were isolated by a modified acid extraction protocol [45]. In briefly, cells and tissues were incubated with extraction buffer (PBS with 0.5% Triton X-100, 1 mM PMSF and 5 mM sodium butyrate) with gentle rotation at 4 °C for 10 min, and nuclei were isolated by centrifugation (1500 g for 10 min at 4°C). 0.4 M HCL was used to resuspend nuclei, followed by incubation on ice for 30 min, soluble histones were isolated by centrifugation (20,000 g for 10 min at 4°C) and quantified using a Bradford protein assay kit (Beyotime).

Proteins were separated using 6-15% SDS-PAGE and transferred onto PVDF membranes (Millipore). Then, the membranes were blocked in 5% skimmed milk (BD Biosciences) in Tris-buffered saline with 0.1% Tween 20 (TBS-T) and incubated with primary antibodies at 4°C overnight. The membranes were then washed in TBS-T for 10 min, three times, and incubated with secondary antibodies at room temperature for 1-2 h. The immunoreactive products were detected by using SuperSignal™ West Pico PLUS Chemiluminescent Substrate (Thermo Fisher Scientific) and Tanon-5200 Chemiluminescent Imaging System (Tanon, Shanghai, China). ImageJ was then used to perform quantitative analysis of western blot results as the ratio of target histone marks or proteins *gray value* to reference proteins *gray value* (H3, β-Actin or β-Tubulin). The detailed information of the primary and secondary antibodies used were summarized in the Additional file 1: Table S4.

For dot blot assay, peptide samples were spotted onto PVDF membranes, the membranes were first blocked at room temperature for 1 h and then washed in TBS-T. Primary and secondary antibodies were incubated at room temperature for 2 h or overnight and 45 min respectively, followed by TBS-T wash for three times and signal detection.

### ATAC-seq and RNA-seq

We used the TruePrep^®^ DNA Library Prep Kit V2 for Illumina (#TD501, Vazyme, Nanjing, China) to construct ATAC-seq libraries in two biological replicates. In brief, a total of 5×10 ^4^ cells (P19 EC cells-derived NSPCs and digested telencephalon cells at E13.5) were lysed in 50 μL lysis buffer (10 mM Tris-HCl pH 7.4, 10 mM NaCl, 3 mM MgCl_2_ and 0.5% (v/v) NP-40) on ice for 10 min. The suspension was then centrifuged at 500 g for 5 min at 4°C, following by addition of 50 μL trans-position reaction mix containing 5×TTBL and TTE Mix V50 to incubate at 37°C for 30 min. After tagmentation, fragmented DNA was purified with 100 μL (2×) VAHTS^®^ DNA clean beads (Vazyme) and amplified for 12 cycles using the following PCR conditions: 72 °C for 3 min; 98 °C for 30 s; and thermocycling at 98 °C for 15 s, 60 °C for 30 s and 72 °C for 30 s; following by 72 °C for 5 min. After PCR amplification, libraries were purified and selected by DNA clean beads. Libraries were then sequenced on an Illumina paired-end platform with 150 bp per read length by Annoroad Gene Technology Co., Ltd (Beijing, China). mRNA-seq was performed in two biological replicates. Briefly, total RNA was extracted using Trizol™ reagent according to the instructions (Thermo Fisher Scientific) and mRNA was purified from total RNA using poly-T oligo-attached magnetic beads. Libraries were then constructed with the NEBNext^®^ Ultra™ RNA Library Prep Kit for Illumina^®^ (NEB) following the manufacturer’s instructions and sequenced on an Illumina paired-end platform with 150 bp per read length by Annoroad Gene Technology Co., Ltd.

### Native ChIP-seq

Native ChIP was carried out in two biological replicates following the published protocol with minor alterations [12]. In brief, cell pellets (P19 EC cells-derived NSPCs and digested telencephalon cells at E13.5 and E16.5) were resuspended in 500 μL swollen buffer (0.3 M sucrose, 60 mM KCl, 15 mM NaCl, 5 mM MgCl_2_, 0.1 mM EDTA, 15 mM Tris-HCl pH 7.5, 5 mM sodium butyrate, and 1×complete proteinase inhibitor). Nuclei were isolated using an equal volume extraction buffer (0.3 M sucrose, 60 mM KCl, 15 mM NaCl, 5 mM MgCl_2_, 0.1 mM EDTA, 15 mM Tris-HCl, pH 7.5, 0.4 % NP-40, 5 mM sodium butyrate and 1×complete proteinase inhibitor) with gentle rotation (30 rpm) at 4 °C for 10 min. The cell suspension was transferred into new tubes containing 500 μL sucrose cushion buffer (0.6 M sucrose, 60 mM KCl, 15 mM NaCl, 5 mM MgCl_2_, 0.1 mM EDTA, 15 mM Tris-HCl pH 7.5, 5 mM sodium butyrate and 1×complete proteinase inhibitor) and centrifuge at 1500 g for 10 min at 4°C. Nuclei were softly resuspended by MNase digestion buffer (0.32 M sucrose, 4 mM MgCl_2_, 1 mM CaCl_2_, 50 mM Tris-HCl pH 7.5, 5 mM sodium butyrate and 1×complete proteinase inhibitor). The amount of chromatin present in the sample was assessed by spectrophotometry in the presence of 0.1% SDS, and chromatin was then digested using Micrococcal nuclease (Takara) (1U per 10 μg chromatin) at 600 rpm in a thermomixer for 5 min at 37°C. The reaction was stopped by adding EDTA to a final concentration of 5 mM and incubated on ice for 5 min, then cleared at 12,000 g for 10 min at 4°C. The supernatant (S1 fraction) was stored at 4°C and the pellet was resuspended in dialysis buffer (0.2 mM EDTA, 1 mM Tris-HCl pH 7.5, 5 mM sodium butyrate), and then dialyse against 1.5 L dialysis buffer by Slide-A-Lyzer™ Dialysis Cassette, 10K MWCO, 3 mL (Thermo Fisher Scientific) overnight at 4°C. The dialysed pellet (S2 fraction) was cleared at 20,000 g for 10 min at 4°C and combined with S1 fraction for immunoprecipitation.

We measured the dsDNA amount of combined S1 and S2 fraction using Qubit^®^ 4.0 fluorometer (Thermo Fisher Scientific) by Equalbit^®^ 1×dsDNA HS Assay Kit (Vazyme) to enable an accurate calculation of the chromatin amount used for immunoprecipitation experiments. Finally, 25 μg chromatin (dsDNA) was immunoprecipitated using 1.25 μg H3K9ac (PTM Biolabs, Hangzhou, China) and H3K9cr (PTM Biolabs), and 150 μg chromatin was immunoprecipitated using 3 μg H3K18la (PTM Biolabs) overnight at 4°C with gentle rotation (20 rpm) in ChIP dilution buffer (1% Triton X-100, 2 mM EDTA, 150 mM NaCl, 50 mM Tri-HCl pH 7.5, 5 mM sodium butyrate and 1× complete proteinase inhibitor). Antibodies-chromatin complexes were then pulled-down using 20 μL Magna ChIP™ protein A+G magnetic beads (Millipore) by incubating with rotation (20 rpm) for 2 h at 4 °C. Beads were washed one time with low salt wash buffer (0.1% SDS, 1% Triton X-100, 2 mM EDTA, 150 mM NaCl and 20 mM Tris-HCl pH 8.1), high salt wash buffer (0.1% SDS, 1% Triton X-100, 2 mM EDTA, 500 mM NaCl and 20 mM Tris-HCl pH 8.1), LiCl wash buffer (250 mM LiCl, 1% NP-40, 10 mM Tris-HCl pH 8.1, 1% sodium deoxycholate and 1 mM EDTA) and two times with TE buffer (1 mM EDTA and 10 mM Tris-HCl pH 8.1) for 10 min each at 4 °C and 30 rpm. Immune complexes were eluted from beads twice for 15 min at 65 °C and 1000 rpm in a thermomixer using 250 μL elution buffer (1% SDS and 100 mM NaHCO_3_).

The eluted immune complexes were then treated with RNase A (100 μg/ml) for 30 min at 37 °C, and followed by Proteinase K (50 μg/ml) for 2 h at 58 °C and 600 rpm in a thermomixer. Then, the IPed DNA was purified using the phenol-chloroform-isoamyl alcohol (25:24:1) method and precipitated with two volumes of 100% ethanol and 10 ug linear acrylamide (Thermo Fisher Scientific) at -20 °C for 2h. Libraries were then constructed with the TruSeq ^®^ ChIP Library Preparation Kit (Illumina) following the manufacturer’s instructions and sequenced on an Illumina paired-end platform with 150 bp per read length by Annoroad Gene Technology Co., Ltd.

### Sequencing data filtering

The original image data obtained after sequencing was converted into the sequence data stored in FASTQ file format by base calling. Raw reads data were filtered by Trimmomatic (v0.36) to remove contaminations and low-quality reads with the parameters “ILLUMINACLIP:Adapter.fa:2:30:10 SLIDINGWINDOW:4:15 LEADING:3 TRAILING:3 MINLEN:36” and quality-controlled using FastQC (v0.11.7) [46, 47].

### ATAC-seq data analysis

The reads of ATAC-seq data were aligned using Bowtie 2 (v2.4.1) to the mouse reference genome with parameters “-X 1000 --mm --local” [48]. Samtools (v1.10) was then used to convert files to bam format and filter reads mapped with parameters “-F 1804 -f 2 -q 30” [49]. After removing PCR duplicates using Mark Duplicates function in Picard (v2.22.0) and mitochondrial reads. MACS2 (v2.2.7.1) was used to call peaks with parameters “--keep-dup all -q 0.01 -f BAMPE” [50].

### ChIP-seq data analysis

High-quality ChIP-seq reads were aligned using Bowtie 2 to the mouse reference genome using default parameters. Then samtools was used to convert files to bam format and filter reads mapped with parameters “-F 1804 -f 2 -q 30”. After removing PCR duplicates using Mark Duplicates function in Picard and mitochondrial reads. MACS2 was used to call peaks with parameters “-q 0.01 -f BAMPE” relative to input samples.

### RNA-seq data analysis

RNA-seq data were quantified using Salmon (v1.1.0) with the parameter “-i -g --gcBias -- validateMappings” [51]. Differential gene expression analysis was conducted using DESeq2 (v1.26.0) and differentially expressed genes were defined by p.adjust < 0.05 and absolute fold change more than 0 [52].

### Peak annotation and gene enrichment analysis

Peak annotation was performed using ChIPseeker (v1.22.1) at gene level and the promoter regions was defined as +/-2 kb of TSS [53]. Genes were considered under regulation of histone marks with at least one peak annotated in their promoter regions or gene body regions. Gene enrichment analysis was performed using clusterProfiler (v3.14.3) with default parameters [54].

### Different binding analysis of ChIP-seq data

For different binding analysis of histone marks in the promoter regions, BEDTools (v2.29.0) “coverage” function was used for read counting within +/-2 kb of TSS and then DESeq2 was used to perform different binding analysis [55], genes with different binding of histone marks were defined by p.adjust < 0.05 and absolute fold change more than 1.5. DiffBind (v2.14.0) and MAnorm (v1.2.0) were adopted to perform different binding analysis of histone marks in the peak regions, and peaks with different binding of histone marks were defined by p.adjust < 0.05 and absolute fold change more than 1.5 [56, 57]. Genes with different binding of histone marks were defined as at least one peak annotated in their promoter regions or gene body regions. For comparison analysis of histone marks intensity across different groups of NSPCs, BEDTools “coverage” function was employed for read counting within both +/-2 kb of TSS (promoter regions) and gene body regions, then the raw read counts were used to calculate reads per kilobase per million mapped reads (RPKM) values representing ChIP signals as follows: [(read-counts) / (region-length in kb)] / (total mapped reads in Mb).

### Sequencing data visualization

Bam files were converted to bigwig files by bamCoverage function of deepTools (v3.4.0) with the paramters “--binSize 50 --normalizeUsing RPKM”, and “computeMatrix”, “plotHeatmap”, “multiBigwigSummary” and “plotCorrelation” functions were used to generate of heatmaps plots [58]. For genome browser representation, data in bigwig files generated by deepTools were visualized using IGV (v2.4.10) [59]. BETA (v1.0.7) “basic” function (--df 0.05 --da 1 -c 0.001) was used for activating and repressive function prediction of peaks with different binding of histone marks [35]. Peaks of replicates were pulled by BEDtools “intersect” function, and bigwig files of replicates were merged by deepTools “bigwigCompare” functions with the parameter “--operation mean”. The mouse reference genome sequence (vM24) and gene annotation (vM24) were downloaded from GENCODE. Summary of ChIP-seq, ATAC-seq and RNA-seq data analysis were provided in the Additional file 1: Table S5.

## Declarations

### Ethics approval and consent to participate

Not applicable.

### Consent for publication

Not applicable.

### Availability of data and materials

The ATAC-seq, ChIP-seq and RNA-seq datasets generated and analysed during the current study are available in NCBI’s GEO and are accessible through series accession number GSE171088 and GSE171832.

### Authors’ contributions

Conceptualization and methodology, S.K.D., and C.M.L.; Investigation, S.K.D., and P.P.L.; Writing-original draft, S.K.D.; Writing-review & editing, S.K.D., C.M.L., and Z.Q.T.; Supervision, C.M.L. All authors have read and approved the manuscript.

## Acknowledgements

Not applicable.

## Supplementary Information

**Additional file 1:** Fig. S1-S10 and Table S4, S5.

**Additional file 2:** Table S1. List of genes with different binding of H3K9ac, H3K9cr or H3K18la in the promoter regions across different developmental stages (E16.5 vs E13.5).

**Additional file 3:** Table S2. List of genes under regulation of H3K9ac, H3K9cr or H3K18la with different binding of these marks across different developmental stages (E16.5 vs E13.5).

**Additional file 4:** Table S3. List of genes under regulation of H3K9ac, H3K9cr or H3K18la with different binding of these marks under HDAC1-3 inhibition (MS-275 vs DMSO).

**Additional file 5:** Uncropped western blotting analysis.

