## Additional file 1 for "Dynamic profiling and functional interpretation of histone lysine crotonylation and lactylation during neural development"

**Content Page**

Fig. S1 Dynamic changes of histone Kcr and Kla in vivo and in cells. (**3**)

Fig. S2 Dot blot assay of antibodies used in this study. (**5**)

Fig. S3 Pearson correlation and cluster analysis of ChIP-seq data. (**6**)

Fig. S4 Multi-omics profiling of H3K9cr and H3K18la in NSPCs. (**8**)

Fig. S5 Enrichment analysis of different expressed genes across developmental stages. (**10**)

Fig. S6 Different binding analysis of histone lysine acylations in the promoter regions. (**11**)

Fig. S7 Different binding analysis of histone lysine acylations in the peak regions. (**13**)

Fig. S8 Genome-wide changes of H3K9cr and H3K18la in vivo. (**15**)

Fig. S9 Genome-wide changes of histone lysine acylations in NSPCs. (**16**)

Fig. S10 Histone lysine acylations regulate gene expression in NSPCs. (**18**)

Table S4. Summary of antibodies used in this study. (**20**)

Table S5. Summary of ChIP-seq, ATAC-seq and RNA-seq data analysis. (**23**)

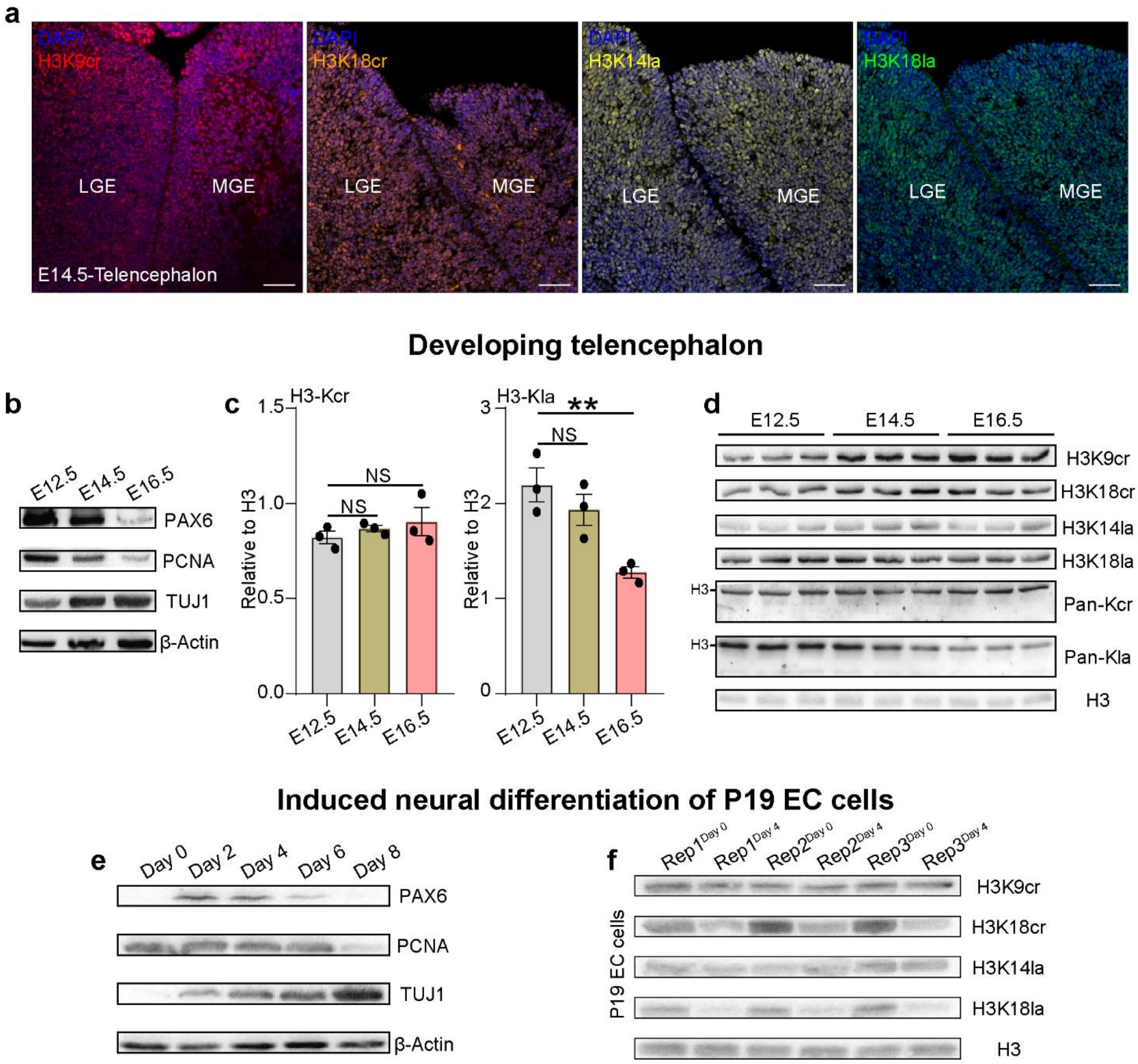


**Fig. S1** Dynamic changes of histone Kcr and Kla in vivo and in cells. **a** Immunostaining of indicated histone marks on mouse forebrain sections at E14.5 (n = 1), LGE: lateral ganglionic eminence and MGE: ganglionic eminence. Scale bar, 50 μm. **b** Western blot assay of PAX6, PCNA and TUJ1 expression in different development stages of telencephalon, which indicated robust neurogenesis (n = 1). **c** Statical analysis for changes of H3-Kcr and H3-Kla levels in the developing telencephalon by western blot assay (n = 3). **d** Original images of western blot assay of indicated histone marks in the developing telencephalon (n = 3). **e** Western blot assay of

PAX6, PCNA and TUJ1 expression across various time points of RA-induced neural differentiation of P19 EC cells, which demonstrated successfully induced neural differentiation of P19 EC cells (n = 1). **f** Original images of western blot assay of indicated histone marks in P19 EC cells-derived day 4 NSPCs and day 8 neurons (n = 3). In (**c**), data were represented as mean ± SEM; NS: no significance; **P< 0.01 by two-tailed unpaired Student’s t-test.

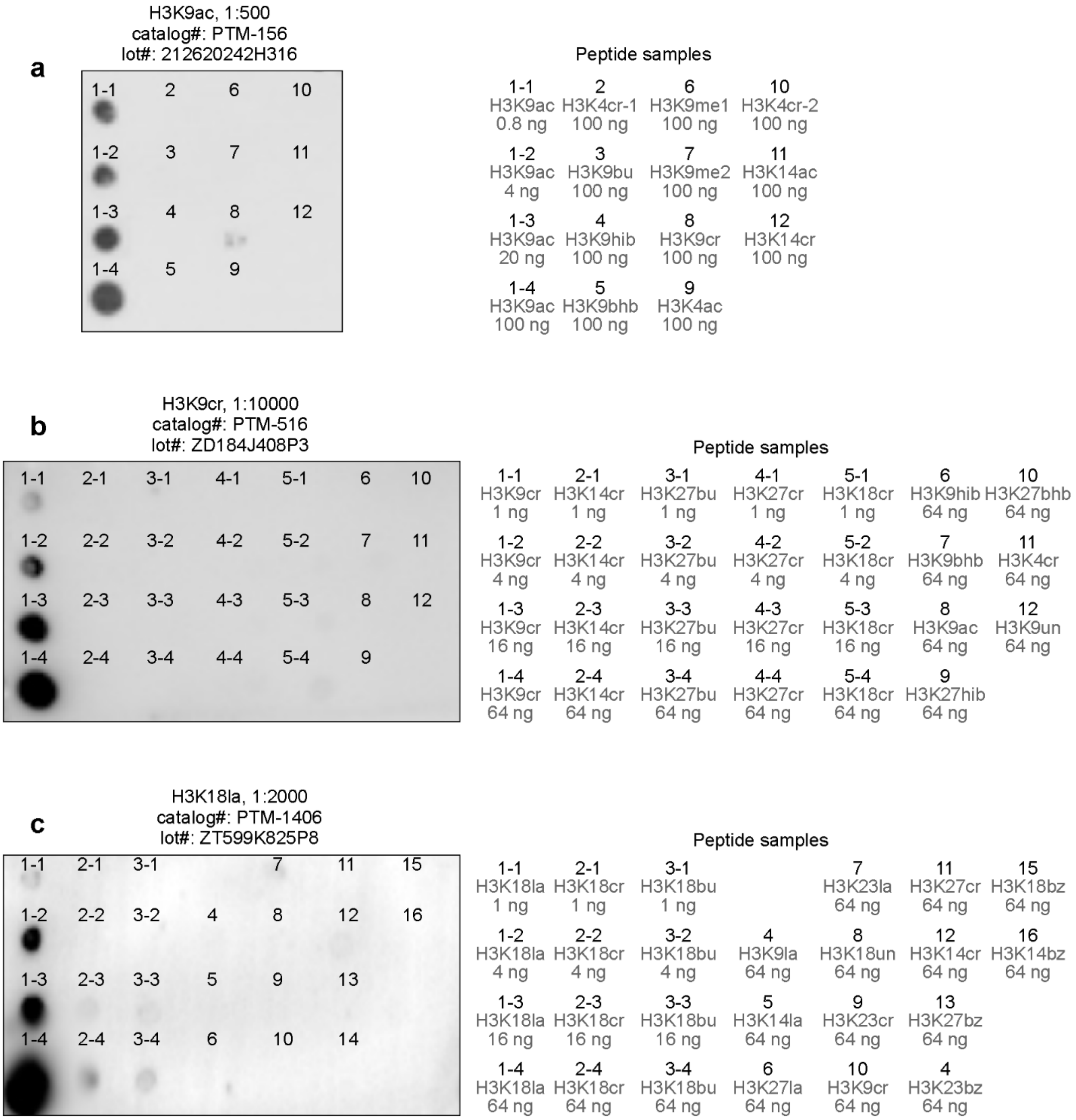


**Fig. S2** Dot blot assay of antibodies used in this study. **a-c** Dot blot assay of H3K9ac (**a**), H3K9cr (**b**) and H3K18la (**c**) antibodies demonstrating its specificity, and the peptide samples used were listed on the right.

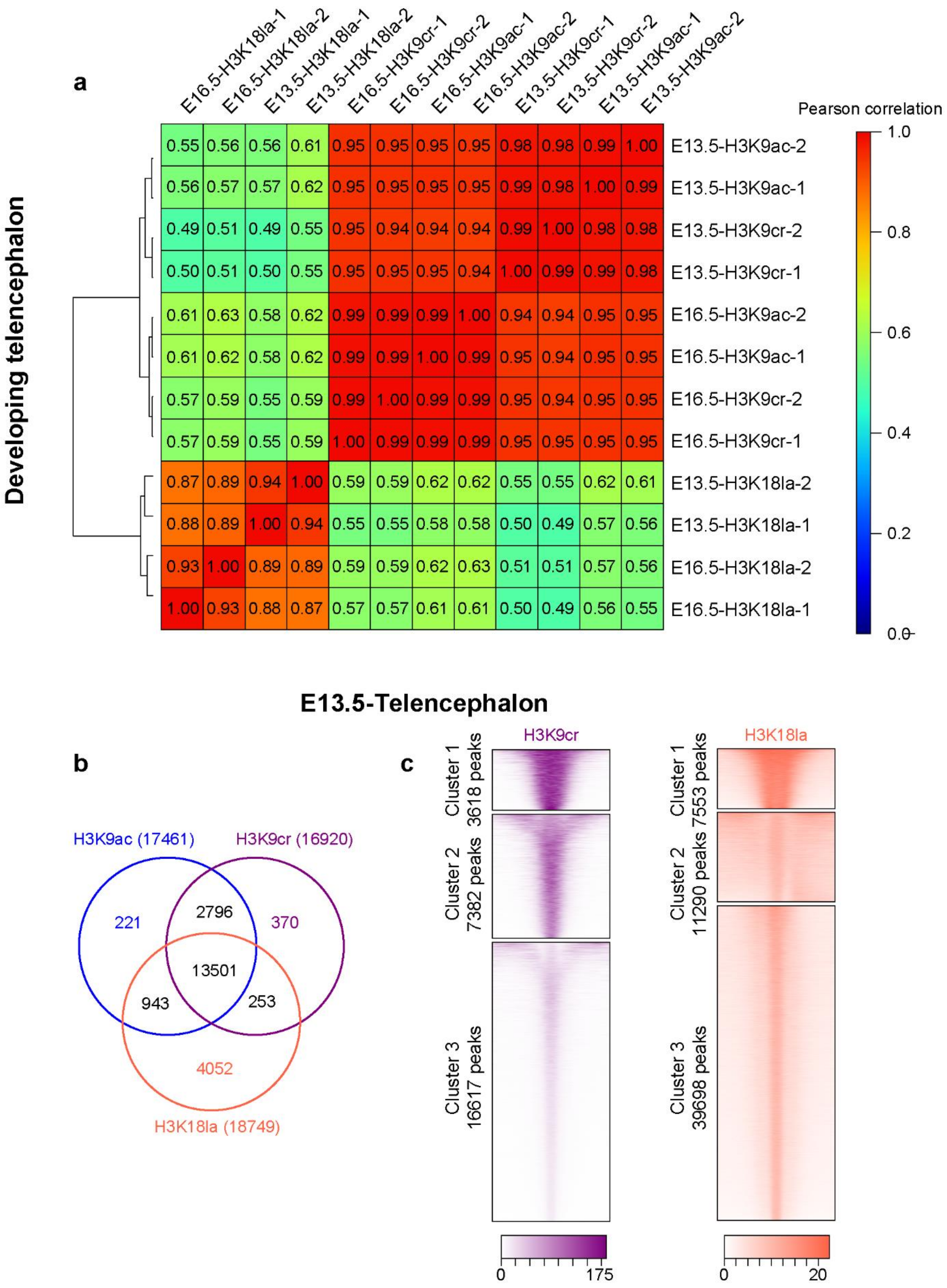


**Fig. S3** Pearson correlation and cluster analysis of ChIP-seq data. **a** The pearson correlation of ChIP-seq data across different developmental stages. **b** Venn diagram of combined comparison of genes under regulation of H3K9ac (17461 genes), H3K9cr (16920 genes) or H3K18la (18749 genes) in the E13.5 telencephalon. **c** Heatmaps at peaks of H3K9cr or H3K18la (scaled by RPKM) plotted within +/- 2.5 kb of the peak center (5’ to 3’) in the E13.5 telencephalon, which were divided to into three clusters and ranked from the highest to the lowest by ChIP signal, the number of peaks in each cluster was labelled on the left.

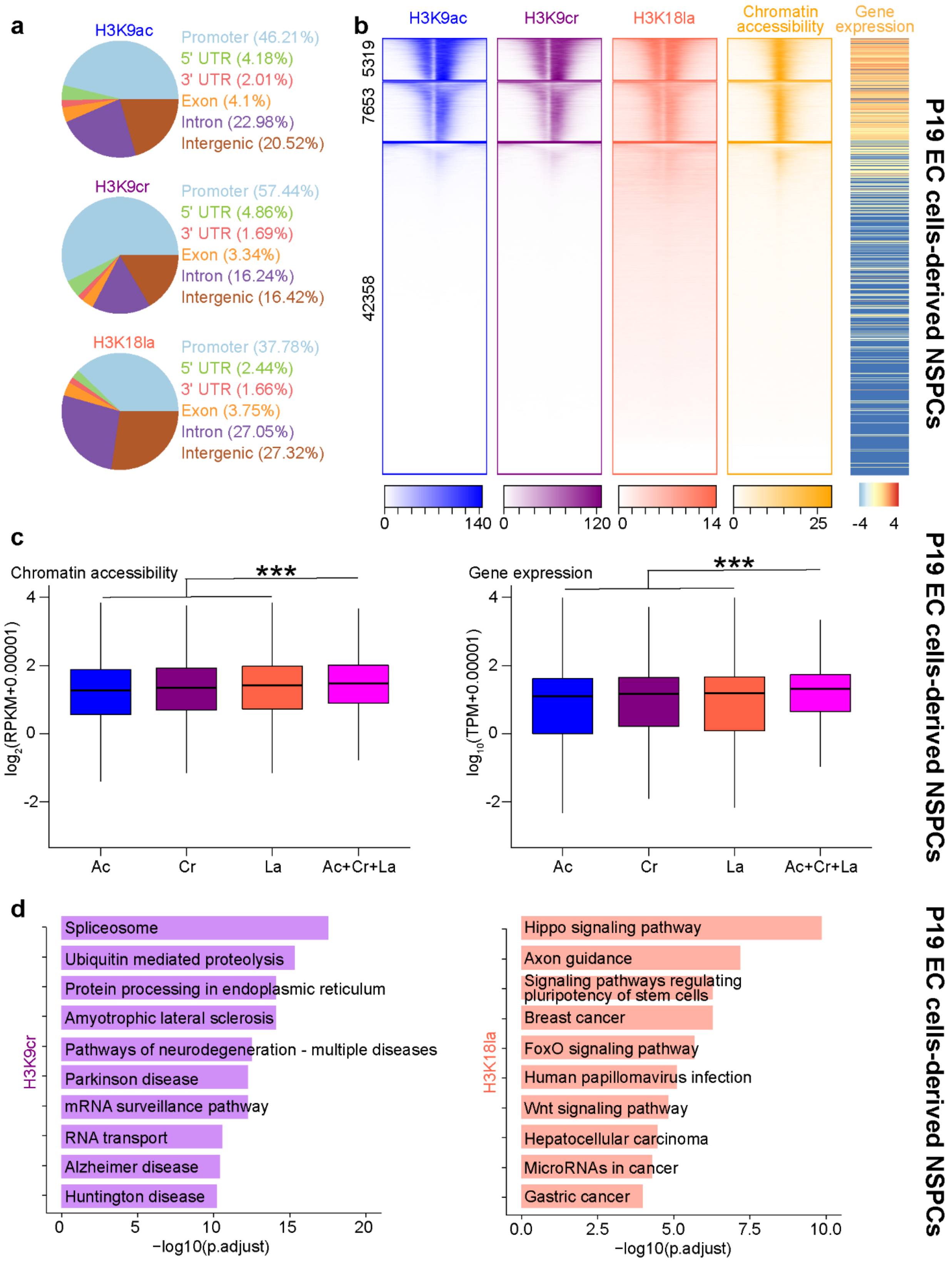


**Fig. S4** Multi-omics profiling of H3K9cr and H3K18la in NSPCs. **a** Pie chart showing distribution of peaks of indicated histone marks at annotated genomic regions in the P19 EC cells-derived day 4 NSPCs. **b** Heatmaps of indicated histone marks intensity (scaled by RPKM), chromatin accessibility levels (scaled by RPKM) plotted within +/- 2.5 kb of the TSS (5' to 3'), and gene expression levels (scaled by $\log_{10}$(TPM+0.00001)) which were divided to into three clusters and ranked from the highest to the lowest by H3K9ac signal in the P19 EC cells-derived day 4 NSPCs, the number of genes in each cluster was labelled on the left. **c** Boxplot showing comparison of chromatin accessibility (left panel) and gene expression (right panel) across different groups of genes in the P19 EC cells-derived day 4 NSPCs. Ac, Cr, La and Ac+Cr+La: genes under regulation of H3K9ac (14450 genes), H3K9cr (13787 genes), H3K18la (12509 genes) or all three marks (10279 genes). **d** KEGG pathway enrichment analysis of genes under regulation of H3K9cr (left panel, 3960 genes) or H3K18la (right panel, 5638 genes) in the P19 EC cells-derived day 4 NSPCs. In (**c**), ***$P < 0.001$ by two-tailed unpaired Student's t-test.

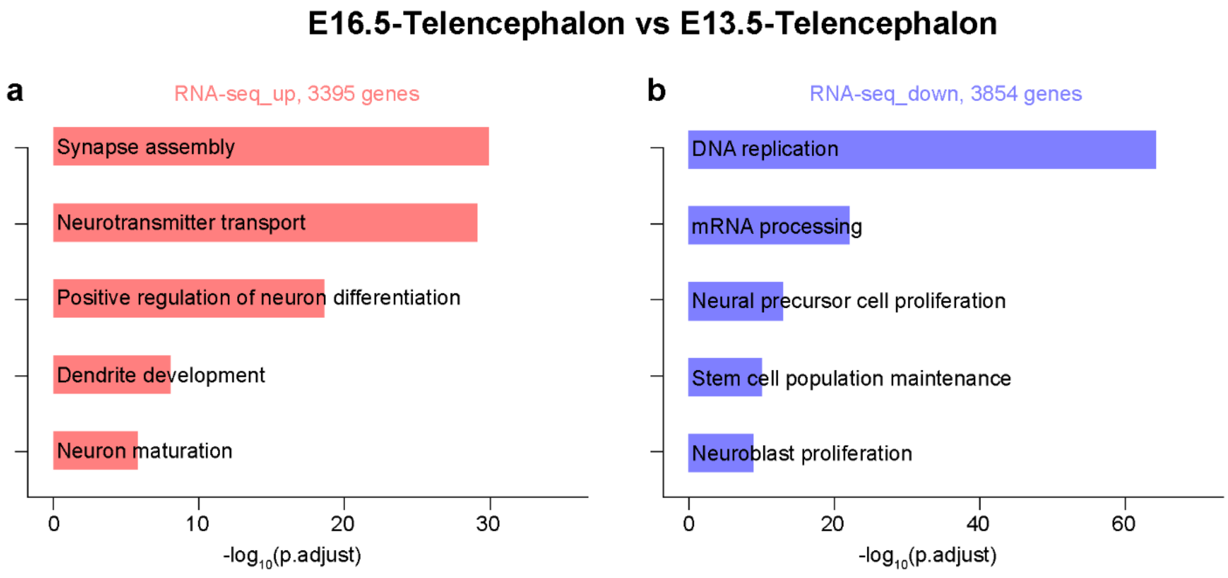


**Fig. S5** Enrichment analysis of different expressed genes across developmental stages. **a, b** GO enrichment analysis (biological process) of up-regulated genes (**a**) or down-regulated genes (**b**) during neural development, which suggested cell fate transition across different stages of neurogenesis.

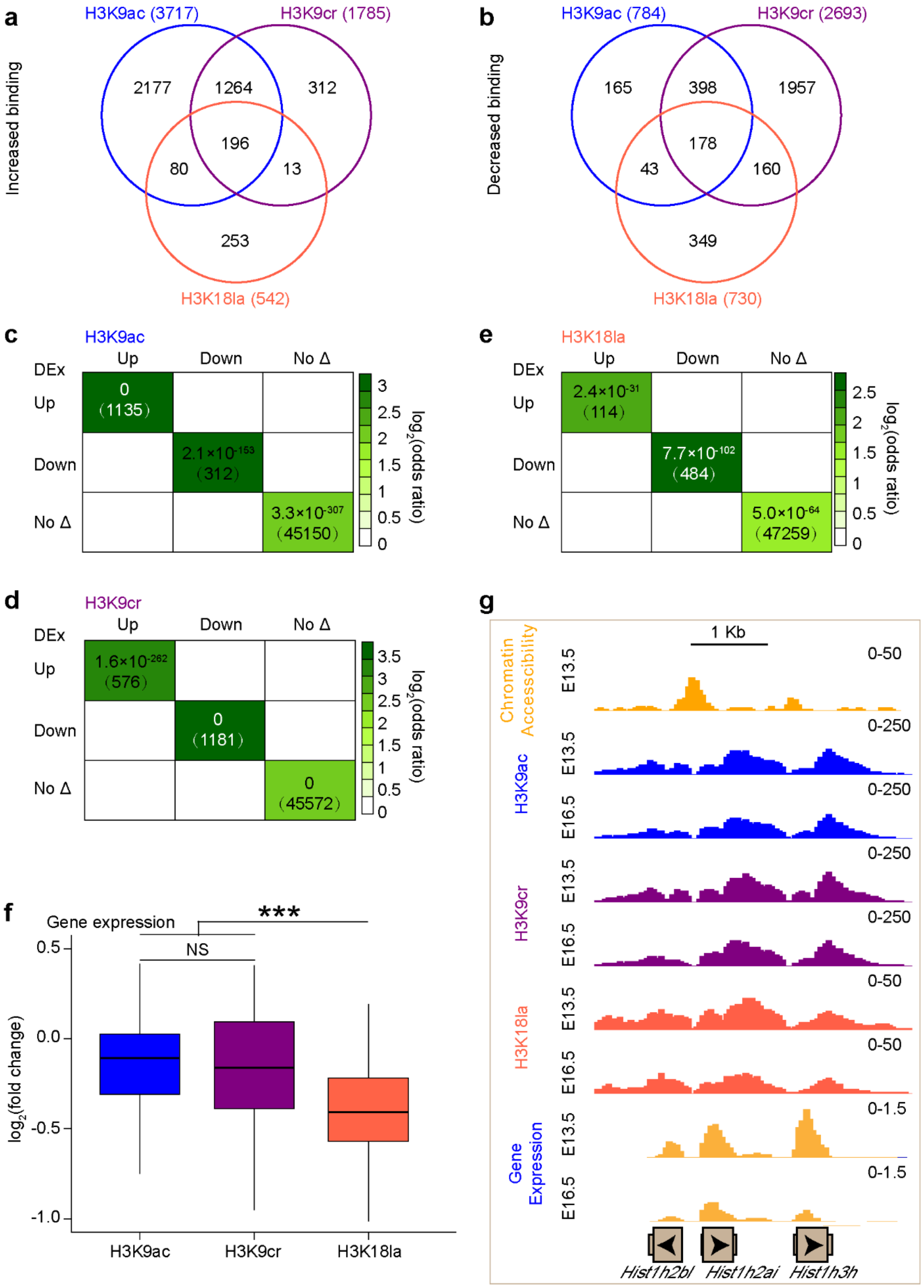


**Fig. S6** Different binding analysis of histone lysine acylations in the promoter regions. **a, b** Venn diagram of a combined comparison of groups of genes with increased (**a**) or decreased (**b**) binding of indicated histone marks in the promoter regions. **c-e** Odds-ratio analysis of overlapping genes displaying differential H3K9ac (**c**), H3K9cr (**d**) or H3K18la (**e**) enrichment versus differential gene expression (DEx), insert numbers indicate respective P-values for associations, with the number of genes overlapping in parentheses. **f** Boxplot showing comparison of changes of H3K9ac, H3K9cr or H3K18la enrichment in the promoter regions of 70 genes encoding histones. **g** Representative genome-browser view at Hist1h2bl, Hist1h2ai, Hist1h3h gene of different sequencing data sets. In (**f**), NS: no significance; ***$P < 0.001$ by two-tailed unpaired Student's t-test.

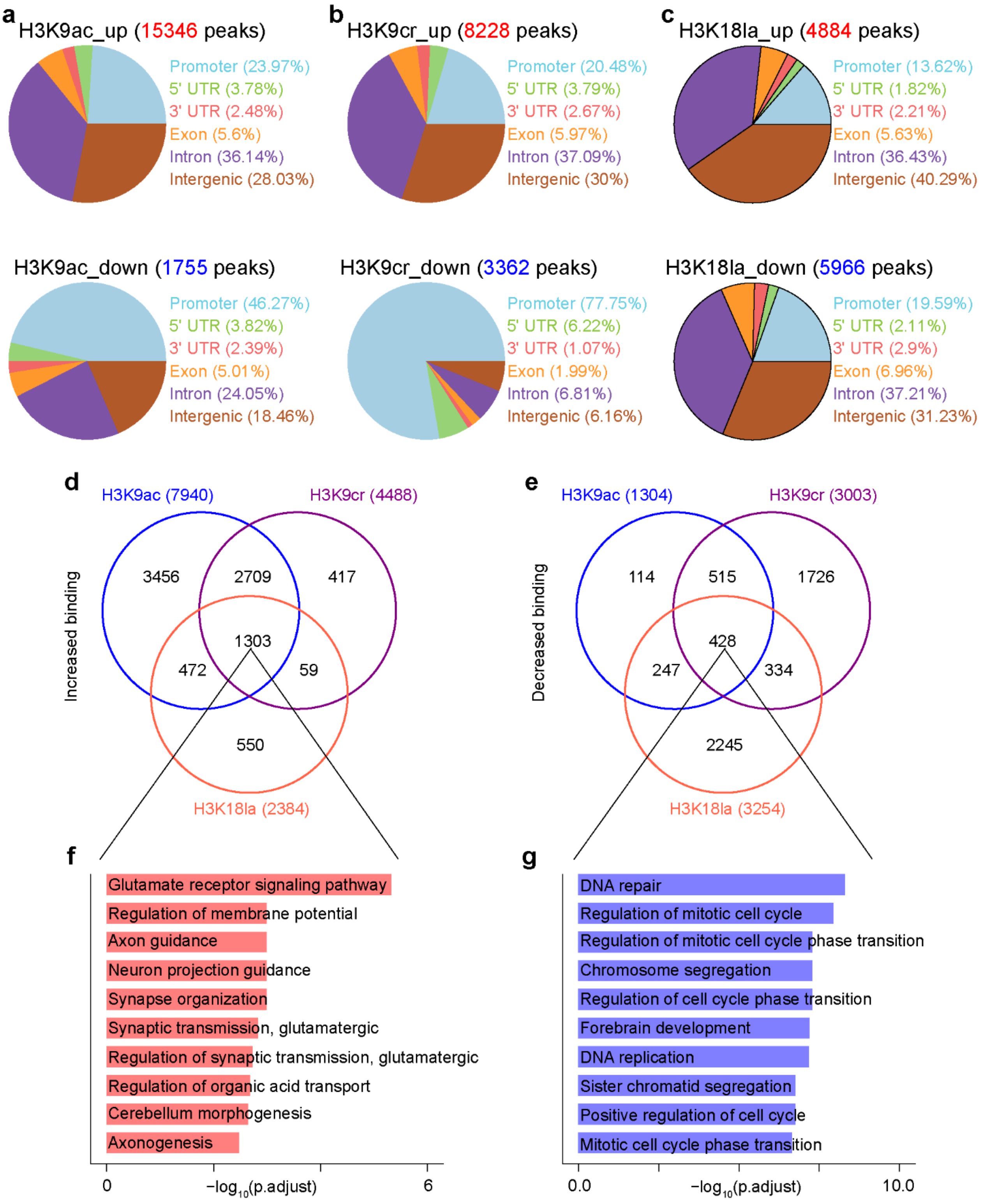


**Fig. S7** Different binding analysis of histone lysine acylations in the peak regions. **a-c** Pie chart showing distribution of peaks with increased binding (top panels) of H3K9ac (**a**), H3K9cr (**b**)

or H3K18la (**c**), as well as decreased binding (bottom panels) of H3K9ac (**a**), H3K9cr (**b**) or H3K18la (**c**) at annotated genomic regions respectively. **d, e** Venn diagram of a combined comparison of groups of genes under regulation of H3K9ac, H3K9cr or H3K18la, with increased (**d**) or decreased (**e**) binding of each mark respectively. **f, g** GO enrichment analysis (biological process) of genes overlapped by increased (**f**) H3K9ac, H3K9cr and H3K18la binding (1303 genes), or decreased (**g**) H3K9ac, H3K9cr and H3K18la binding (428 genes), which suggested that these histone marks cooperatively regulated cell fate transition during neural development.

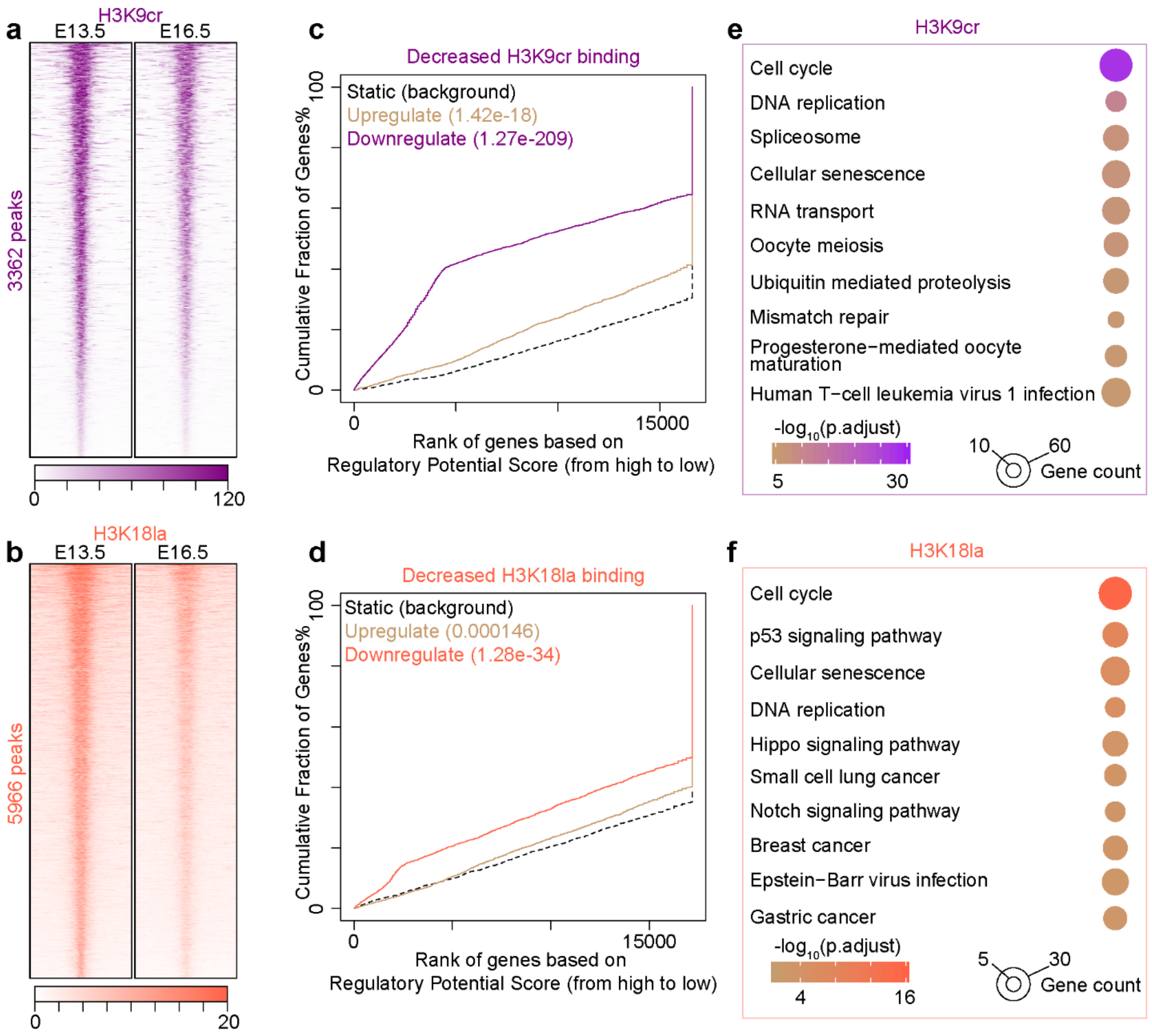


**Fig. S8** Genome-wide changes of H3K9cr and H3K18la in vivo. **a, b** Heatmaps at peaks with decreased H3K9cr (**a**) or H3K18la (**b**) binding (scaled by RPKM) plotted within +/- 5 kb of the peak center (5’ to 3’), the number of peaks was labelled on the left. **c, d** BETA plot of combined computational analysis of ChIP-seq and RNA-seq data using peaks with decreased H3K9cr (**c**) or H3K18cr (**d**) binding. **e, f** KEGG pathway enrichment analysis of down-regulated genes with decreased H3K9cr (**e**) or H3K18la binding (**f**), there were 923 and 1342 genes in (**e**) and (**f**) respectively.

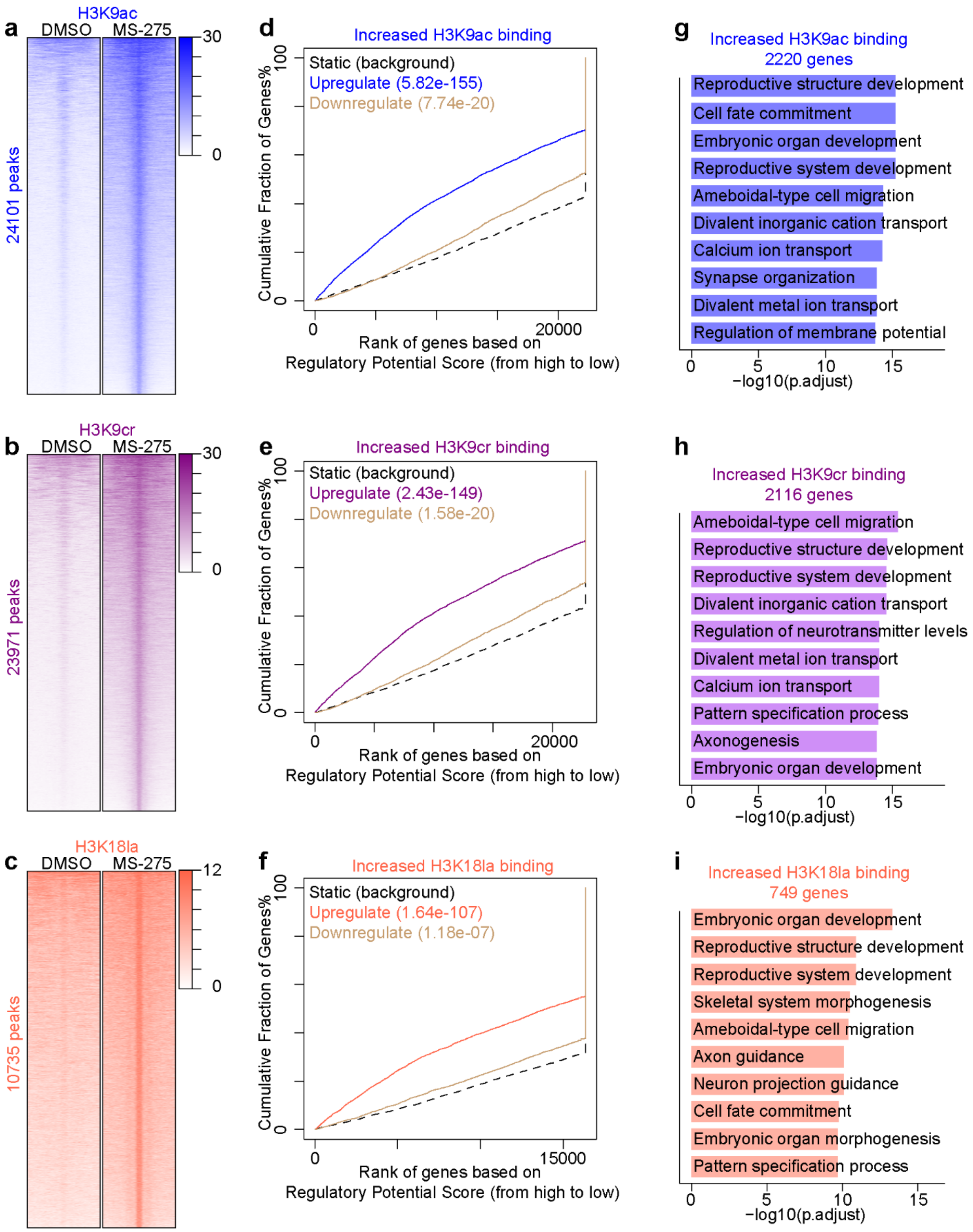


**Fig. S9** Genome-wide changes of histone lysine acylations in NSPCs. **a-c** Heatmaps at peaks with increased H3K9ac (**a**), H3K9cr (**b**) or H3K18la (**c**) binding (scaled by RPKM) plotted within +/- 5 kb of the peak center (5' to 3'), the number of peaks was labelled on the left. **d-f** BETA plot of combined computational analysis of ChIP-seq and RNA-seq data using peaks with increased H3K9ac (**a**), H3K9cr (**b**) or H3K18la (**c**) binding. **g-i** GO enrichment analysis (biological process) of up-regulated genes with increased H3K9ac (**g**), H3K9cr (**h**) or H3K18la (**i**) binding, there were 2220, 2116 and 749 genes in (**g**), (**h**) and (**i**) respectively.

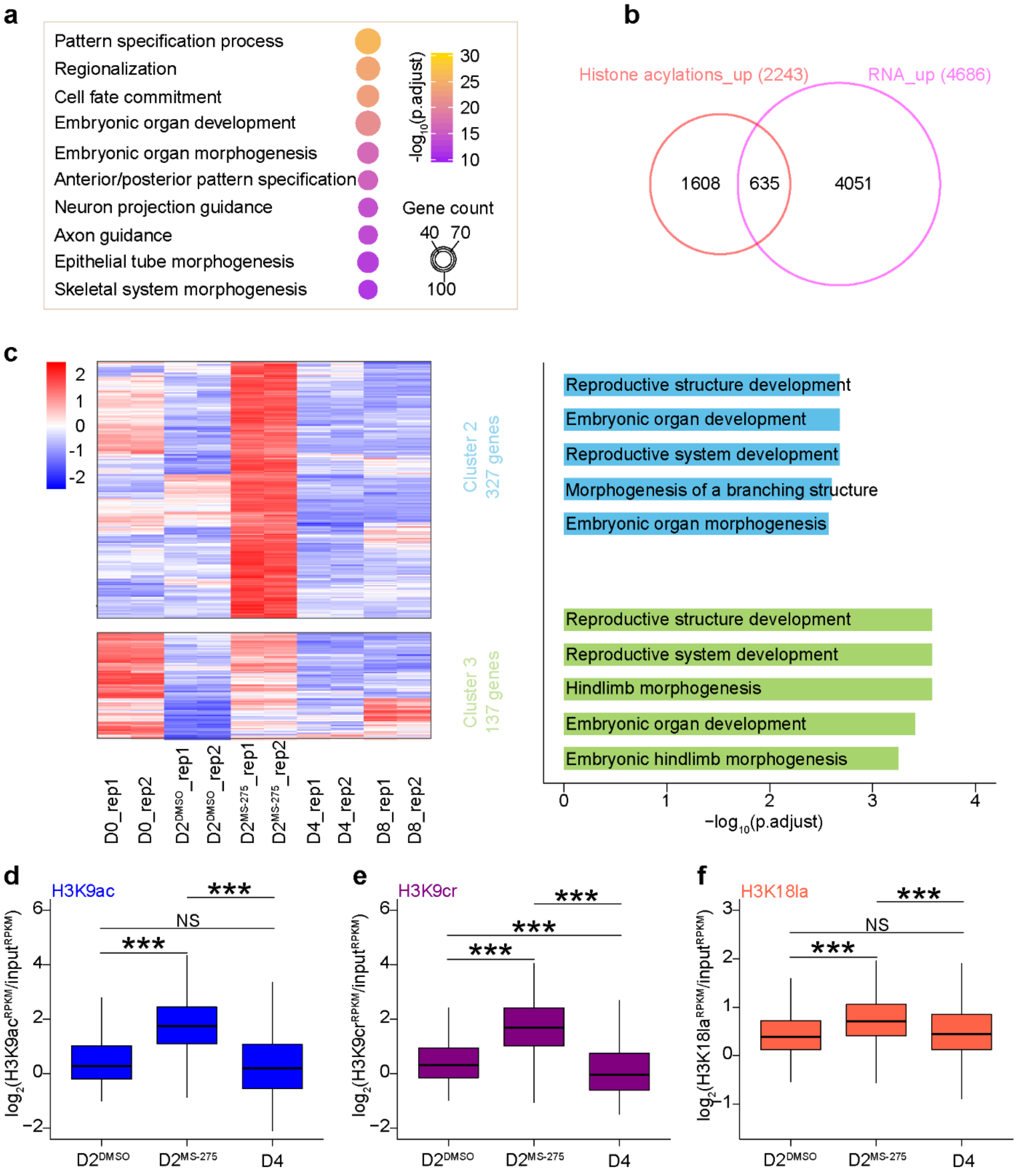


**Fig. S10** Histone lysine acylations regulate gene expression in NSPCs. **a** GO enrichment analysis (biological process) of genes with increased binding of all three marks (H3K9ac, H3K9cr and H3K18la) after MS-275 treatment in P19 EC cells-derived NSPCs (2243 genes). **b** Venn diagram of a combined comparison of groups of genes with increased binding of all three marks (H3K9ac, H3K9cr and H3K18la) and up-regulated expression after MS-275 treatment in P19 EC cells-derived NSPCs. **c** Heatmap showing dynamic changes of up-regulated genes (DESeq2 vsd-normalized RNA-seq counts and scaled by row) with increased binding of all three marks (H3K9ac, H3K9cr and H3K18la) after MS-275 treatment during neural differentiation process of P19 EC cells, and these genes were divided into three clusters (see also Fig. 6d) according their expression patterns (left panel), with the results of GO enrichment analysis (biological process) of genes in each cluster on right panel. **d-f** Boxplot showing comparison of changes of H3K9ac (**d**), H3K9cr (**e**) or H3K18 (**f**) levels across different groups of NSPCs at genes in cluster 2 (327 genes) defined in (**c**), which indicated that genes in cluster 2 got increased binding of these histone marks in MS-275 treated NSPCs at Day2 ($D2^{MS\text{-}275}$) comparing with that in DMSO treated control group ($D2^{DMSO}$), but the increased trends were not monitored during normal neural differentiation (D4 (NSPCs at Day4) vs $D2^{DMSO}$). In (**d-f**), NS: no significance; ***P< 0.001 by two-tailed unpaired Student's t-test.

<table>
<tr><th colspan="5">Table S4. Summary of antibodies used in this study.</th></tr>
<tr><th>Antibody</th><th>Purpose</th><th>Company</th><th>Cat.Number</th><th>Dilution</th></tr>
<tr><td>anti-β-Actin</td><td>Immunoblot</td><td>EASYBIO, Beijing, China</td><td>BE0033</td><td>1:2000-1:5000</td></tr>
<tr><td>anti-H3</td><td>Immunoblot</td><td>EASYBIO</td><td>BE7004</td><td>1:2000-1:5000</td></tr>
<tr><td>anti-HDAC1</td><td>Immunoblot</td><td>Sangon Biotech, Shanghai, China</td><td>D260479</td><td>1:1000</td></tr>
<tr><td>anti-HDAC2</td><td>Immunoblot</td><td>Sangon Biotech</td><td>D155199</td><td>1:1000</td></tr>
<tr><td>anti-HDAC3</td><td>Immunoblot</td><td>Sangon Biotech</td><td>D260481</td><td>1:1000</td></tr>
<tr><td rowspan="2">anti-H3K9ac</td><td>ChIP-seq</td><td rowspan="2">PTM Biolabs, Hangzhou, China</td><td rowspan="2">PTM-156</td><td>1.25 μg/ChIP-seq</td></tr>
<tr><td>Immunoblot</td><td>1:1000</td></tr>
<tr><td rowspan="3">anti-H3K9cr</td><td>ChIP-seq</td><td rowspan="3">PTM Biolabs</td><td rowspan="3">PTM-516</td><td>1.25 μg/ChIP-seq</td></tr>
<tr><td>Immunoblot</td><td>1:1000</td></tr>
<tr><td>Immunostaining</td><td>1:1000</td></tr>
<tr><td>anti-H3K18cr</td><td>Immunoblot</td><td>PTM Biolabs</td><td>PTM-517</td><td>1:2000</td></tr>
</table>

| | | | | |
|---|---|---|---|---|
| | Immunostaining | | | 1:2000 |
| anti-H3K14la | Immunoblot | PTM Biolabs | PTM-1414 | 1:2000 |
| | Immunostaining | | | 1:2000 |
| anti-H3K18la | ChIP-seq | PTM Biolabs | PTM-1406 | 3 μg/ChIP-seq |
| | Immunoblot | | | 1:2000 |
| | Immunostaining | | | 1:2000 |
| anti-H3K27ac | Immunoblot | Abcam | ab4729 | 1:2000 |
| anti-Nestin | Immunostaining | Aves Labs | NES | 1:500 |
| anti-Pan-Kcr | Immunoblot | PTM Biolabs | PTM-501 | 1:1000 |
| anti-Pan-Kla | Immunoblot | PTM Biolabs | PTM-1401 | 1:1000 |
| anti-PAX6 | Immunoblot | Invitrogen | 14-9914 | 1:1000 |
| anti-PCNA | Immunoblot | Cell Signaling Technology | 13110 | 1:1000 |
| anti RNA pol II | Immunoblot | Active Motif | 39497 | 1:2000 |

| | | | | |
|---|---|---|---|---|
| anti RNA pol II CTD repeat phospho S2 | Immunoblot | Abcam | ab193468 | 1:2000 |
| anti RNA pol II CTD repeat phospho S5 | Immunoblot | Abcam | ab193467 | 1:2000 |
| anti-β-Tubulin | Immunoblot | EASYBIO | BE3312 | 1:2000-1:5000 |
| anti-TUJ1 | Immunoblot | BioLegend | 801202 | 1:1000 |
| | Immunostaining | | | 1:1000 |
| donkey anti-rabbit Alexa Fluor 568 | Immunostaining | Thermo Fisher Scientific | A10042 | 1:500 |
| goat anti-chicken Alexa Fluor 594 | Immunostaining | Thermo Fisher Scientific | A11042 | 1:500 |
| goat anti-mouse Alexa Fluor 594 | Immunostaining | Thermo Fisher Scientific | A11032 | 1:500 |
| goat anti-rabbit Alexa Fluor 488 | Immunostaining | Thermo Fisher Scientific | A11034 | 1:500 |
| goat anti-mouse HRP | Immunoblot | EASYBIO | BE0102 | 1:2000-1:5000 |
| goat anti-rabbit HRP | Immunoblot | EASYBIO | BE0101 | 1:2000-1:5000 |

| Table S5. Summary of ChIP-seq, ATAC-seq and RNA-seq data analysis. | | | | | | |
|---|---|---|---|---|---|---|
| ChIP-seq | | | | | | |
| Sample name | Replicate | Total read pairs | Uniquely alignment rate (%) | Overall alignment rate (%) | Total peaks | Regulated genes |
| E13.5-Telencephalon-H3K9ac | 1 | 53724013 | 65.26 | 98.54 | 37319 | 18574 |
| E13.5-Telencephalon-H3K9ac | 2 | 51131802 | 59.39 | 98.25 | 34532 | 18128 |
| E13.5-Telencephalon-H3K9cr | 1 | 46817521 | 61.41 | 97.89 | 32567 | 18115 |
| E13.5-Telencephalon-H3K9cr | 2 | 36679840 | 55.41 | 95.42 | 28968 | 17266 |
| E13.5-Telencephalon-H3K18la | 1 | 46396342 | 77.85 | 98.93 | 73369 | 20519 |
| E13.5-Telencephalon-H3K18la | 2 | 39864161 | 76.50 | 98.95 | 69157 | 20532 |
| E16.5-Telencephalon-H3K9ac | 1 | 45849859 | 67.10 | 98.77 | 49652 | 20996 |
| E16.5-Telencephalon-H3K9ac | 2 | 55471433 | 65.94 | 98.57 | 50668 | 21138 |
| E16.5-Telencephalon-H3K9cr | 1 | 44637264 | 69.87 | 97.33 | 41074 | 19925 |
| E16.5-Telencephalon-H3K9cr | 2 | 46975520 | 73.09 | 98.62 | 42304 | 20141 |
| E16.5-Telencephalon-H3K18la | 1 | 45649297 | 72.71 | 98.61 | 62073 | 18771 |
| E16.5-Telencephalon-H3K18la | 2 | 45469142 | 81.91 | 98.69 | 76037 | 20889 |

| | | | | | | |
|---|---|---|---|---|---|---|
| E13.5-Telencephalon-input | 1 | 52415013 | 61.65 | 98.84 | NA | |
| E13.5-Telencephalon-input | 2 | 50313387 | 61.33 | 98.83 | | |
| E16.5-Telencephalon-input | 1 | 41469169 | 60.83 | 98.83 | | |
| E16.5-Telencephalon-input | 2 | 42552642 | 61.38 | 98.83 | | |
| P19-D4-NSPCs-H3K9ac | 1 | 44385449 | 77.62 | 98.81 | 38703 | 19084 |
| P19-D4-NSPCs-H3K9ac | 2 | 46237774 | 78.24 | 98.72 | 37578 | 18881 |
| P19-D4-NSPCs-H3K9cr | 1 | 43443207 | 71.42 | 97.71 | 29280 | 17196 |
| P19-D4-NSPCs-H3K9cr | 2 | 43683550 | 70.49 | 98.06 | 28707 | 17184 |
| P19-D4-NSPCs-H3K18la | 1 | 43944019 | 67.43 | 98.43 | 54865 | 18212 |
| P19-D4-NSPCs-H3K18la | 2 | 54241452 | 70.70 | 98.46 | 51057 | 17951 |
| P19-D4-NSPCs-input | 1 | 47291008 | 66.42 | 98.56 | NA | |
| P19-D4-NSPCs-input | 2 | 46914207 | 64.76 | 98.57 | | |
| DMSO_P19-Day 2-H3K9ac | 1 | 46696262 | 67.41 | 97.65 | 26555 | 16495 |
| DMSO_P19-Day 2-H3K9cr | 1 | 47324833 | 70.31 | 97.37 | 24874 | 16069 |
| DMSO_P19-Day 2-H3K18la | 1 | 41266344 | 65.67 | 98.57 | 36382 | 16778 |

| | | | | | | |
|---|---|---|---|---|---|---|
| MS-275_P19-Day 2-H3K9ac | 1 | 39077351 | 72.25 | 98.46 | 84418 | 23736 |
| MS-275_P19-Day 2-H3K9cr | 1 | 61457579 | 62.95 | 97.91 | 70748 | 22251 |
| MS-275_P19-Day 2-H3K18la | 1 | 39838816 | 70.04 | 98.54 | 31577 | 9831 |
| DMSO_P19-Day 2-input | 1 | 53034910 | 56.35 | 98.46 | NA | |
| MS-275_P19-Day 2-input | 1 | 47774132 | 55.9 | 98.04 | | |

| **ATAC-seq** | | | | | |
|---|---|---|---|---|---|
| **Sample name** | **Replicate** | **Total read pairs** | **Uniquely aligned reads (%)** | **Total aligned reads (%)** | **Total peaks** |
| E13.5-Telencephalon | 1 | 32490761 | 31.78 | 61.84 | 26423 |
| E13.5-Telencephalon | 2 | 37379579 | 30.10 | 57.64 | 28207 |
| P19-D4-NSPCs | 1 | 42851384 | 39.35 | 95.77 | 28399 |
| P19-D4-NSPCs | 2 | 37858578 | 40.21 | 96.66 | 24420 |

| **RNA-seq** | | | |
|---|---|---|---|
| **Sample name** | **Replicate** | **Total read pairs** | **Mapping rate (%)** |
| E13.5-Telencephalon | 1 | 20944513 | 81.45 |

| | | | |
|---|---|---|---|
| E13.5-Telencephalon | 2 | 22718954 | 81.37 |
| E16.5-Telencephalon | 1 | 22498548 | 77.93 |
| E16.5-Telencephalon | 2 | 23373976 | 77.11 |
| P19-D0-ECCs | 1 | 23547414 | 82.95 |
| P19-D0-ECCs | 2 | 22022756 | 82.73 |
| P19-D4-NSPCs | 1 | 23954541 | 81.42 |
| P19-D4-NSPCs | 2 | 22657641 | 81.78 |
| P19-D8-Neurons | 1 | 23707915 | 81.89 |
| P19-D8-Neurons | 2 | 22849732 | 81.32 |
| DMSO_P19-Day 2 | 1 | 23809406 | 82.26 |
| DMSO_P19-Day 2 | 2 | 22205497 | 81.58 |
| MS-275_P19-Day 2 | 1 | 23019633 | 81.91 |
| MS-275_P19-Day 2 | 2 | 23174365 | 81.14 |
