## Supplementary figures and images for "Dynamic profiling and functional interpretation of histone lysine crotonylation and lactylation during neural development"

### Additional file 5

**Fig. 4**

**Fig. 4a**

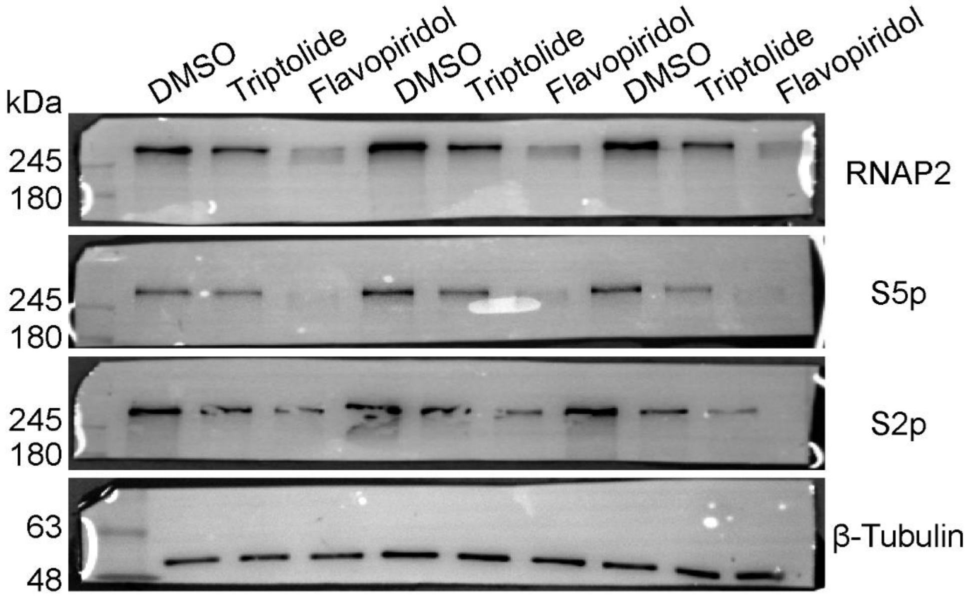


**Fig. 4c**

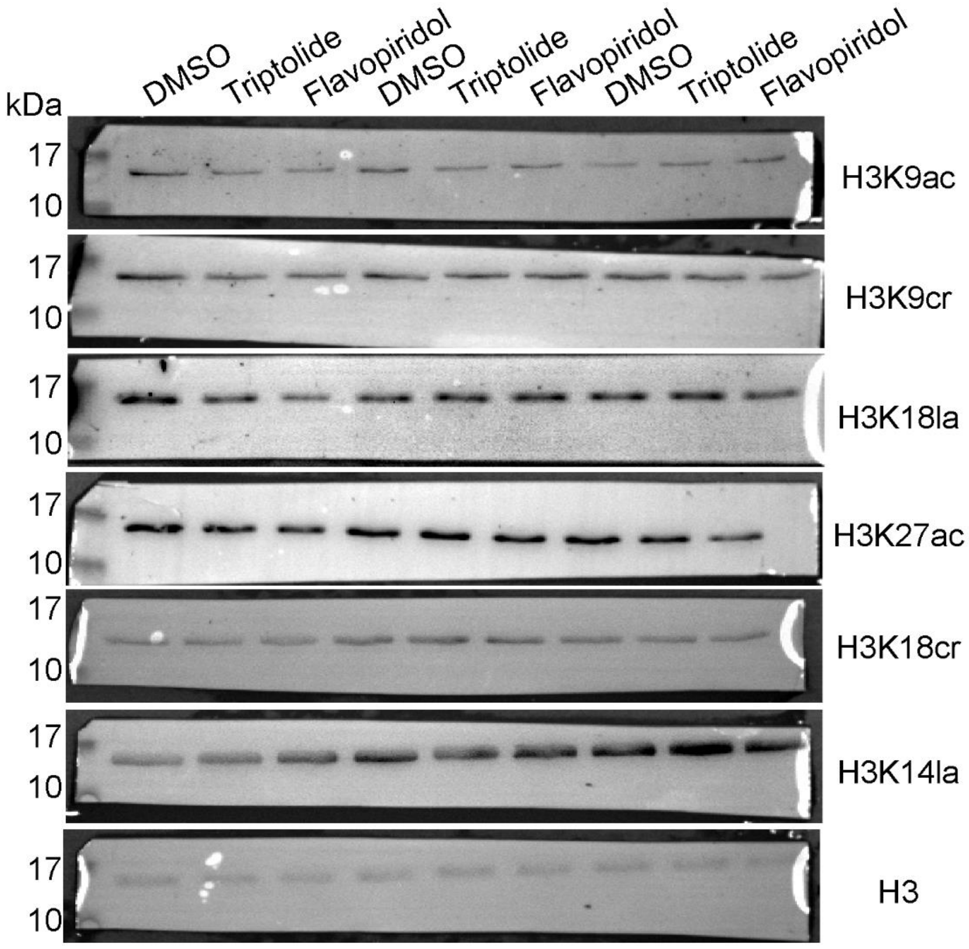

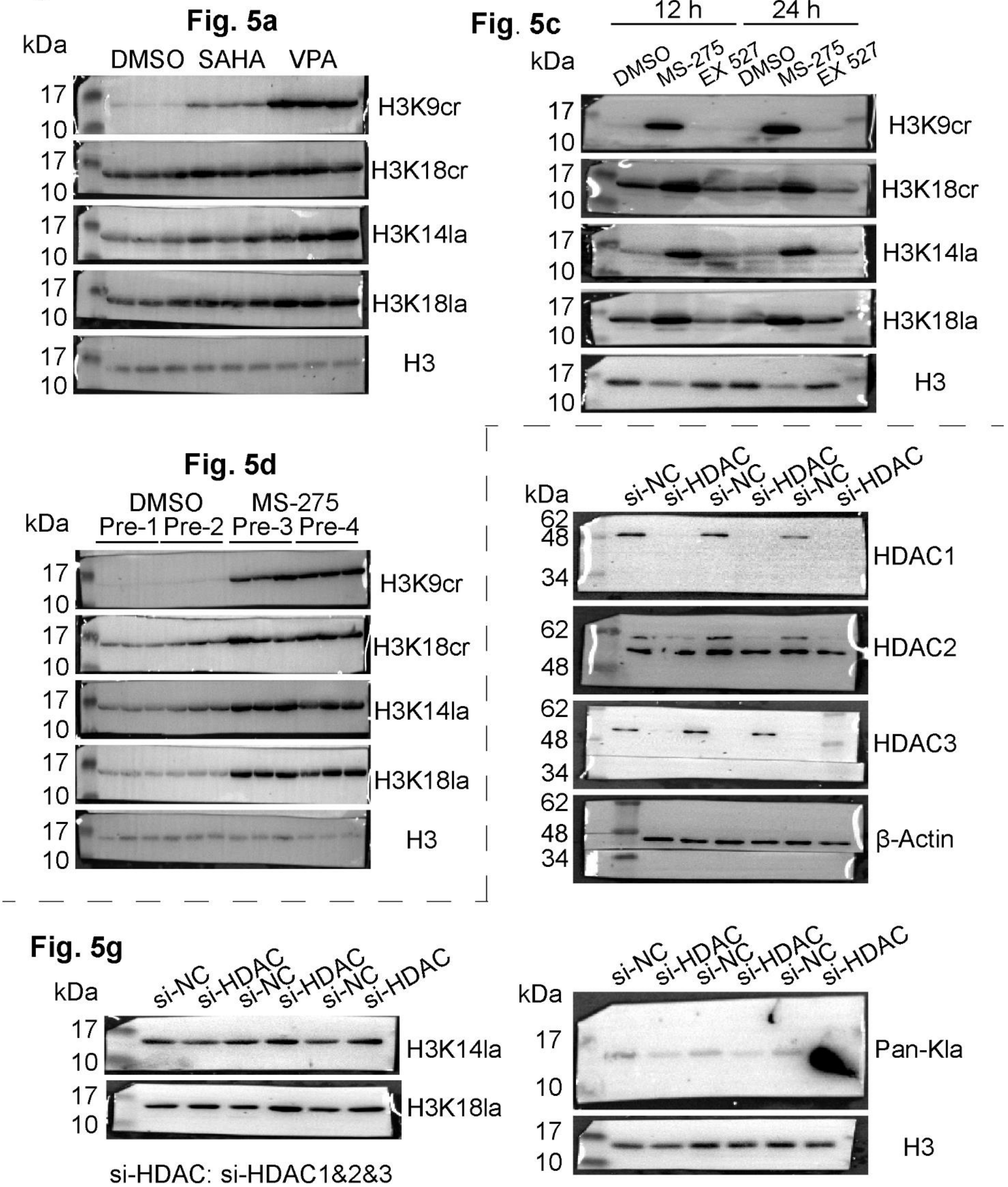

#### Additional file 1: Fig. S1

##### Fig. S1b

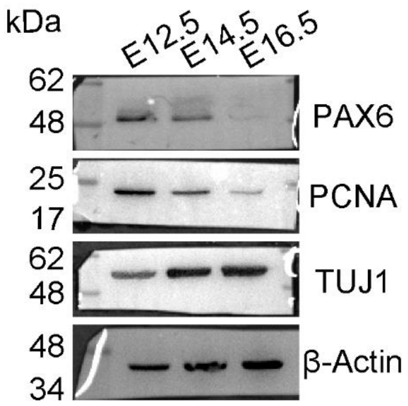


##### Fig. S1d

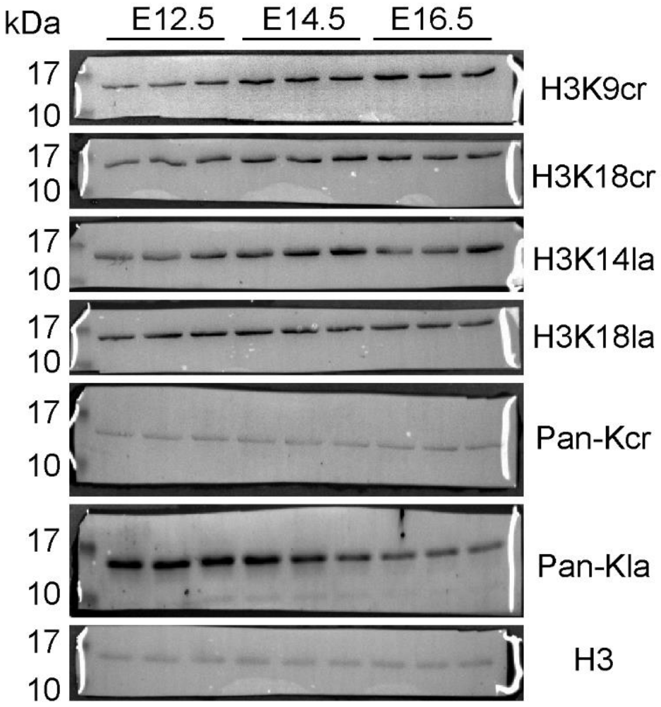


##### Fig. S1e

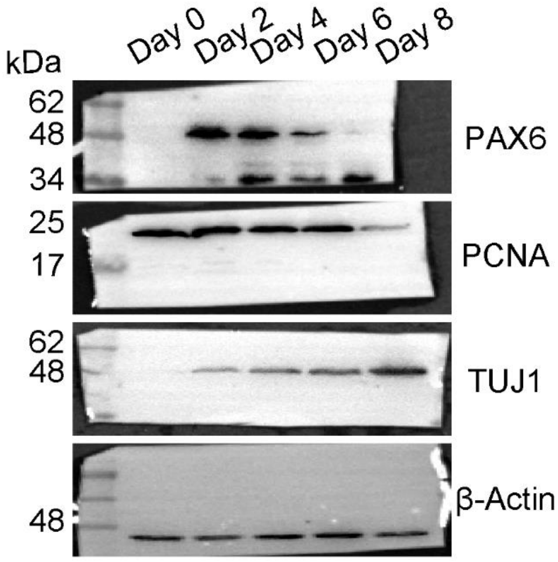


##### Fig. S1f

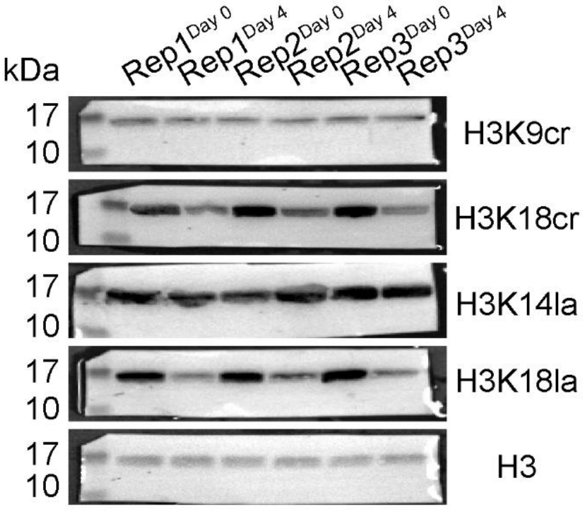
